# Allosteric effects of the coupling cation in melibiose transporter MelB

**DOI:** 10.1101/2025.07.10.664195

**Authors:** Parameswaran Hariharan, Yuqi Shi, Bakhtiiari Amirhossein, Ruibin Liang, Rosa Viner, Lan Guan

**Affiliations:** Department of Cell Physiology and Molecular Biophysics, Center for Membrane Protein Research, School of Medicine, Texas Tech University Health Sciences Center, Lubbock, TX, USA; Thermo Fisher Scientific, San Jose, CA, USA; Department of Chemistry and Biochemistry, Texas Tech University, Lubbock, TX, USA

**Author notes:** To whom correspondence should be addressed: Lan Guan.

**Keywords:** Allostery, symport, structural dynamics, sugar recognition, x-ray crystallography, HDX-MS, Molecular Dynamics simulations

## Abstract

The major facilitator superfamily (MFS) transporters play significant roles in human health and disease. *Salmonella enterica* serovar Typhimurium melibiose permease (MelB_St_), which catalyzes the symport of galactosides with Na^+^, H^+^, or Li^+^, is a prototype of this important transporter superfamily. We have published the structures of the inward- and outward-facing conformations of MelB_St_ with galactoside or Na^+^ bound, determined the binding thermodynamic cycle, and proposed that positive cooperativity between the two co-transported solutes plays a key role in the symport mechanism of MelB_St_. The molecular basis for this core mechanism remains unclear. In this study, we determined the molecular basis for this core symport mechanism through analyzing the structural dynamics of MelB_St_ and effects induced by melibiose, Na^+^, or both using hydrogen-deuterium exchange mass spectrometry (HDX-MS). We also refined the specific determinants for the sugar recognition in both protein and galactoside molecules by solving the crystal structures of a uniporter D59C MelB_St_ bound to melibiose and other sugars, and identified a critical water molecule as part of sugar recognition. Our integrated studies from structure, HDX-MS, and molecular dynamics simulations support the conclusion that sugar-binding affinity is directly correlated with protein dynamics. The binding of the coupling cation at a remote site functions as an allosteric activator to restrain the conformational flexibility of dynamic residues in the sugar-binding site and in the cytoplasmic gating salt-bridge network, thereby increasing sugar-binding affinity allosterically. This study provides a molecular-level schematic of the fundamental symport mechanism via positive cooperativity, which may serve as a general mechanism for cation-coupled symporters.

## Introduction

The Solute Carrier (SLC) family of transporters encompasses diverse superfamilies of membrane proteins with various protein folds and employing different transport mechanisms to facilitate the translocation of a wide range of solutes across cell membranes (1). The largest superfamily of SLC transporters is the major facilitator superfamily (MFS)(2), which includes a significant number of cation-coupled secondary active transporters. MFS transporters are responsible for the uptake of a broad spectrum of solutes across cell membranes, playing crucial roles in physiology, pathology, and pharmacokinetics, and are emerging as drug targets (3, 4). Recent rapid advancements in membrane protein research have greatly enhanced our understanding of protein conformation and mechanisms(5–8); however, critical details remain lacking. For example, in cation-coupled symport, it is still unclear how the coupling cations facilitate the binding, translocation, and accumulation of the primary substrate.

For MFS secondary active transporters, most members use H^+^ as the coupling cation, and a few members use Na^+^, such as the Na^+^-coupled essential lipid transporter (MFSD2A) that is expressed in the major organ barriers, including the blood-brain barrier or blood-retina barrier (9–11). The Na^+^-coupled melibiose transporter of *Salmonella enterica* serovar Typhimurium (MelB_St_), which is a well-characterized representative for the Na^+^-coupled MFS transporters, catalyzes the symport of a galactopyranoside with Na^+^, H^+^, or Li^+^, and is a valuable model system for studying cation-coupled transport mechanisms (12–22). Two major conformations of MelB_St_ have been determined: an outward-facing conformation at the apo or galactoside-bound states(20, 22) and an Na^+^-bound inward-facing conformation (21). The primary substrate-specificity determinant pocket and the cation-specificity determinant pocket have been structurally and functionally characterized (19–23). All three coupling cations compete for the same binding pocket, and the transport stoichiometry is 1 galactoside: 1 cation (Na^+^, H^+^, or Li^+^). Binding of the primary and coupling substrates is positively cooperative; the sugar affinity depends on the cation identity, with the cooperativity numbers (fold of increase in affinity) being 8, 5, or 2 for Na^+^, Li^+^, and H^+^, respectively (19, 20). In addition to being sensitive to the binding of the coupling cation and its identity, notably, the sugar-binding affinity is also dependent on MelB_St_ conformation (21). By trapping MelB_St_ in an inward-facing state using the inward-facing conformation-specific binder nanobody-725 (Nb725), both experimental sugar-binding assays and cryoEM structural analysis support that the sugar-binding pocket at inward-facing conformation is at a low-affinity state (21, 23). Remarkably, Na^+^ binding to the inward-facing conformation remains unchanged (21, 23). These results provide experimental evidence supporting the previously proposed stepped-binding kinetic model for melibiose/Na^+^ symport, in which Na^+^ binds first and is released after the sugar release on the opposite surface (15, 18, 24, 25).

Positive cooperativity of substrate binding has been proposed to be the key symport mechanism in MelB_St_, but the molecular basis for this critical mechanism remains unclear. The structures indicate that the bound sugar and Na^+^ have no direct contact, while the two binding pockets are in close proximity (20, 21). The minimum free-energy landscape for sugar translocation, which was simulated based on the structures of the outward- and inward-facing conformations as two starting points (26), suggests that the Na^+^ contribution to the binding free energy of sugar by direct contact is negligible. Cooperativity occurs through allosteric coupling, likely via electrostatic interactions.

In this study, we further analyzed the sugar-binding site by improving the crystal structure resolution of α-NPG bound at 2.60 Å, which uncovered an important water molecule in the binding site. We also confirmed the sugar specificity determinants in both sugar and MelB_St_ by determining the crystal structures in complex with three other α-sugar substrates—melibiose, raffinose, or α-methyl galactoside (α-MG)—that vary in the number of sugar units but all contain an α-galactosyl group. We then examined the structural dynamics of the entire MelB with hydrogen-deuterium exchange coupled to mass spectrometry (HDX-MS) and the effects of melibiose, Na^+^, or both on MelB_St_. HDX-MS is particularly attractive for dynamic systems, as the provided information yields beyond the stable events within the dynamic repertoire of transporter proteins. By monitoring the HDX rate across a time window, it can provide predictive modeling for overall protein folds (27, 28). The resolution of structural information provided by this technique is typically on the peptide level, though. We also performed MD simulations to analyze the side-chain dynamics. Collectively, all results support the notion that the sugar-binding affinity of MelB_St_ is coupled to protein structural dynamics and conformational transitions between inward- and outward-facing states, and the coupling cation in this symporter functions as an allosteric activator.

## Results

### Sugar transport and binding affinity measurements

The sugar analog α-NPG has been determined as a substrate for MelB_Ec_ by measuring *p*-nitrophenol production from the cells that expressed both MelB_Ec_ and α-galactosidase(29). The same assay was modified to determine the α-NPG translocation mediated by MelB_St_ in DW2 cells. Melibiose at 1 mM was added to induce α-galactosidase expression during cell growth, and the washed cells were applied to measure the time-course of *p*-nitrophenol release into the media upon adding 1 mM α-NPG into the induced cells. The detection was for the *p*-nitrophenol, which was resulted from α-NPG transport and hydrolysis. The results showed that both WT MelB_St_ and D59C uniport mutant mediated the translocation of α-NPG (**Fig. 1a**) at this downhill mode of transport.

**Fig 1.**
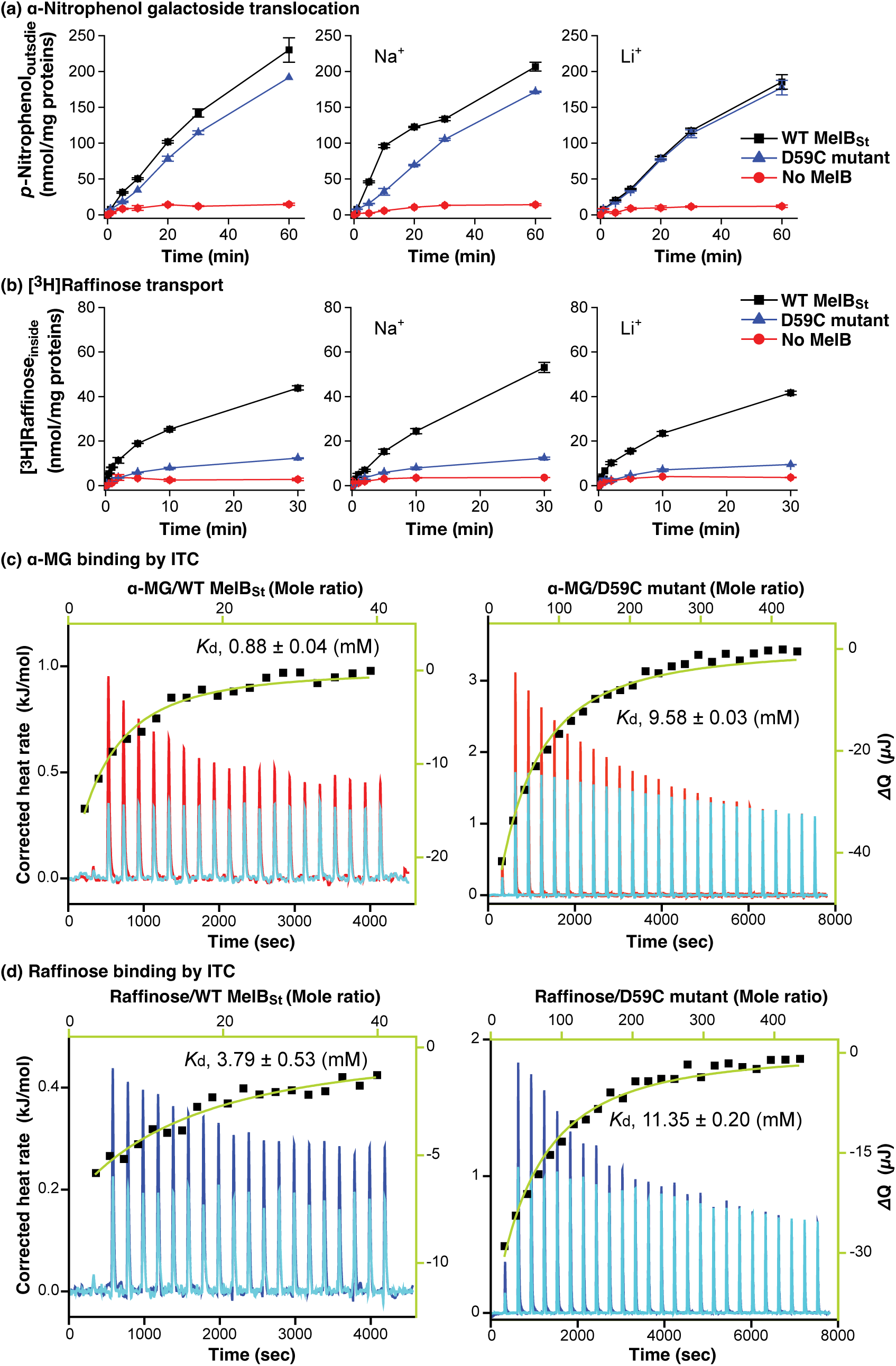
Functional characterizations. **(a)** α-NPG down-hill transport. *E. coli* DW2 cells with the induced α-galactosidase by melibiose in the absence or presence of WT MelB_St_ or D59C mutant were washed before incubating with 0.5 mM *p*-nitrophenyl-α-D-galactoside at 30 °C in the absence or presence of 20 mM NaCl or 20 mM LiCl as described in Methods. The cell-aliquots at given time points at 0, 1, 5, 10, 20, 30, and 60 min were quenched with 0.3 M Na_2_CO_3_, followed by centrifugation to remove the cells. The *p*-nitrophenol in supernatant released from the cells was measured at *A*_405_ nm. **(b)** [^3^H]Raffinose active transport. The *E. coli* DW2 cells in the absence or presence of MelB_St_ with no α-galactosidase induction were used for the active transport of [^3^H]raffinose at 1 mM (specific activity, 10 mCi/mmol) at 23 °C in the absence or presence of 50 mM NaCl or LiCl. The cellular uptake time course measurements at 0, 0.08, 0.17, 0.5, 1, 2, 5, 10, and 30 min were carried out by a dilution and fast-filtration method. **(c & d)** ITC measurement of α-MG (c) or raffinose (d). ITC measurements were performed at 25 °C under similar buffer conditions: 20 mM Tris-HCl, pH 7.5, 100 mM NaCl, 10% glycerol, and 0.035% UDM detergent. For each experiment, 80 µM of the purified WT MelB_St_ or D59C mutant was placed in the reaction cell, and methyl α-D-galactoside (α-MG) or raffinose at 10 mM (against the WT) or 100 mM (against D59C) from the syringe are incrementally titrated to generate the thermograms. The curve fitting is performed with a one-site independent-binding model included in the NanoAnalyze software (version 3.7.5). The thermograms were plotted as baseline-corrected heat rate (µJ/sec; left axis) vs. time (bottom axis) for the titrant to MelB_St_ (red for α-MG and blue for raffinose) or to buffer (light blue). The heat change Δ*Q* (µJ; filled black symbol) was plotted against the mole ratio of the sugar to WT MelB_St_ (top/right axes in green). **(c)** α-MG. (d) Raffinose.

To determine raffinose transport activity, [^3^H]raffinose transport in the *E. coli* DW2 strain was carried out in the absence of Na^+^ and Li^+^ or the presence of Na^+^ or Li^+^ (**Fig. 1b**). The results showed that raffinose, a trisaccharide formed from melibiose and fructose, is also a substrate and transported by MelB_St_.

Previously, our ITC studies in the presence of Na^+^ revealed that the binding affinity, *K*_d_ values, of the WT and D59C MelB_St_ for melibiose were 1.25 mM ± 0.05 mM and 4.96 ± 0.11 mM, respectively, and for α-NPG were 16.46 ± 0.21 µM or 11.97 ± 0.09 µM, respectively (19, 20, 25, 30). The same assay was utilized to determine the α-MG and raffinose binding in the presence of Na^+^ (**Fig. 1**). By injecting the α-MG or raffinose in a buffer-matched solution into the sample cell containing the WT MelB_St_ or the D59C uniporter mutant in the presence of Na^+^, the isotherm curve fitted well, yielding *K*_d_ values of 0.88 ± 0.04 mM or 3.79 ± 0.53 mM, for α-MG or raffinose, respectively. Raffinose, which has more sugar units, exhibits a poor binding affinity to MelB_St_.

### Outward-facing crystal structures of MelB_St_ complexed with varied sugar substrates

The uniporter D59C MelB_St_ mutant exhibits greater thermostability, making it a valuable tool for structural analysis of sugar binding(20). Here we report four crystal structures of D59C MelB_St_ with the endogenous sugar substrate melibiose, and two other α-galactosides, α-methyl galactoside (α-MG) with a single sugar unit or raffinose with three sugar units, as well as α-NPG at an improved resolution. The structure statistics were presented in **Table S1**. All structures adopt a virtually identical outward-facing conformation with RMSD values of less than 0.4 (**Fig. S1**). This typical MFS-fold transporter with 12 transmembrane helices is organized into two six-helix domains linked by the middle loop between helices VI and VII (Loop_6-7_). There are three cytoplasmic helices at the loop_6-7_, loop_8-9_, and the C-terminal tail or lid (**Fig. 2**), named ICH1-3, respectively; and ICH1 and ICH2 run parallel to the membrane bilayer. In all structures, one sugar molecule is bound in the middle of the protein and sandwiched by both N- and C-terminal domains as described previously(20). As indicated by the sliced surface, the binding residues located within the cytoplasmic leaflet of both domains define the edge of the inner barrier, which prevents the sugar from passing across the transporter into the cytoplasm. On the periplasmic side, the open vestibule connects the solvent to the binding pocket. In the α-NPG -bound structure, as reported in PDB ID 8FRH (D59C apo structure) or ID 8FQ9 (D55C mutant with DDMB(22), a PEG molecule was modeled to the density near Arg296 along with helix IX, which could indicate a potential lipid-binding site (**Fig. S1**).

**Fig. 2.**
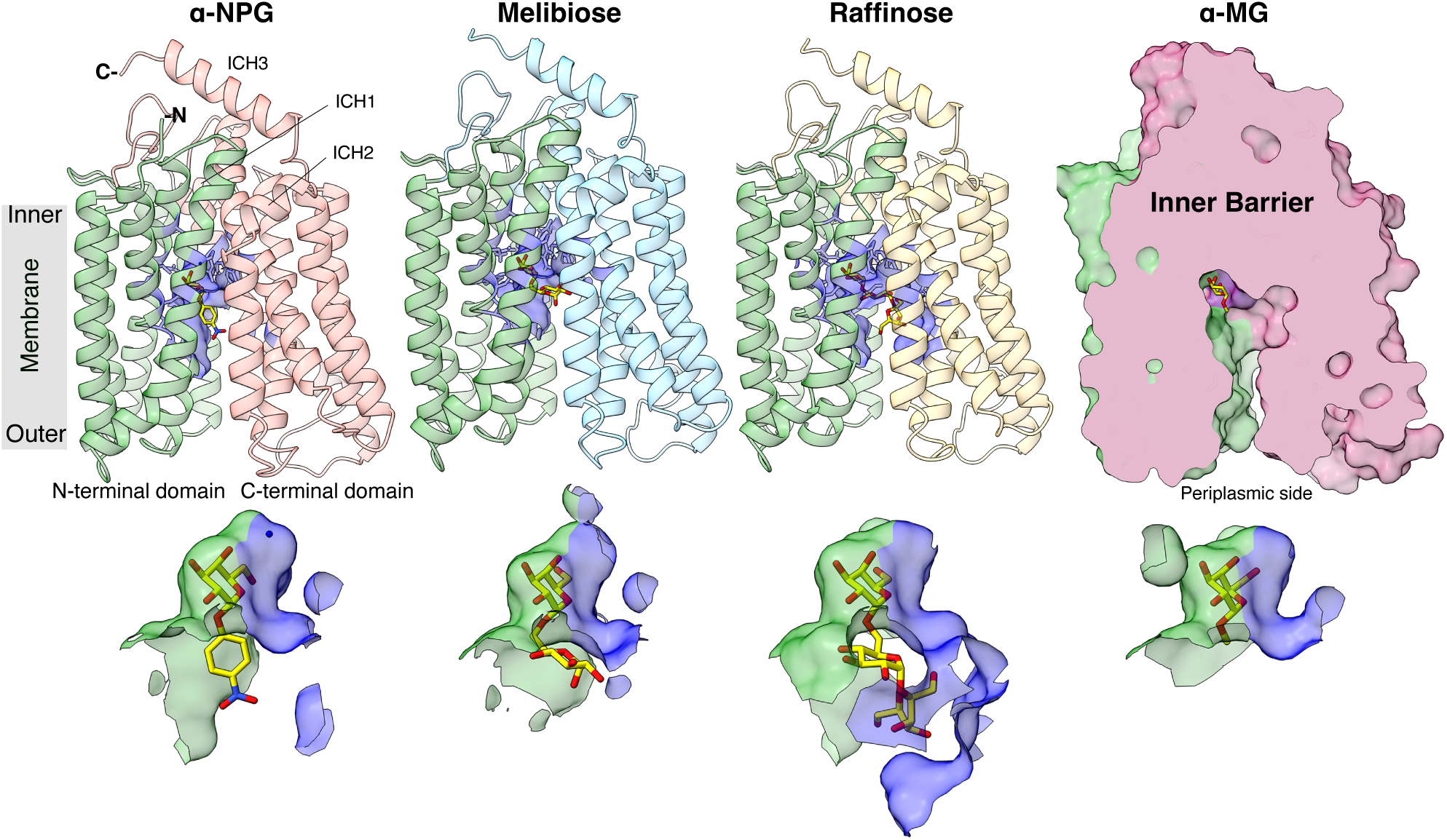
Crystal structures of D59C MelB_St_ in complex with α-sugars substrates. ***Upper row***: Cartoon representation of the structures of D59C MelB_St_ bound with α-NPG, melibiose (α-disaccharide), and raffinose (α-trisaccharide), respectively, along with a surface presentation of D59C MelB_St_ complexed with α-methyl galactoside (α-MG). All structures were oriented with the cytoplasmic side on top and the N-terminal domain (colored green) on the left. Each sugar molecule is colored yellow. The blue sticks and surfaces indicate residues within 5 Å of the sugar molecules. ***Lower row***: Sugar-binding pocket. Residues from the N-terminal and C-terminal domains were shown in surface representation, and colored in green and blue, respectively.

### α-NPG-binding structure refined to a resolution of 2.60 Å

The improved resolution, from 3.01 Å to 2.60 Å, provided a better-resolved density map for the bound α-NPG molecule, which further supported the originally assigned pose, as demonstrated by the stereo view (**Fig. 3**) (20). Thus, the binding pocket is formed by 14 residues on five helices, including N-terminal residues, as labeled in black including helices I (Lys18, Asp19, Ile22, and Tyr26), IV (Tyr120, Asp124, and Tyr128), and V (Arg149 and Ala152), and the C-terminal residues labeled in blue including helices X (Trp342) and XI (Gln372, Thr373, Val376, and Lys377). A water molecule (Wat 1) was modeled to a positive density, which located at hydrogen-bonding distances from the C4-OH and C6-OH on the galactopyranosyl ring and the Thr373 at helix XI, and also surrounded by Gln372 at helix XI and Asp124 at helix IV, as shown in the stereo-view of a 2Fo-Fc electron density map (**Fig. 3**). Another three water molecules participated the cytoplasmic gating salt-bridge network between the both bundles (**Fig. S2a, b**). Water molecules Wat 2 and Wat 3 interacted with the charged pair Arg295 (helix IX) and Asp351 (helix XI), respectively, and Wat 4 interacted with Lys138 from loop_4-5_, which also formed a salt-bridge interaction with Glu142 in this functionally important gating area (**Fig. S2b**) (31).

**Fig. 3.**
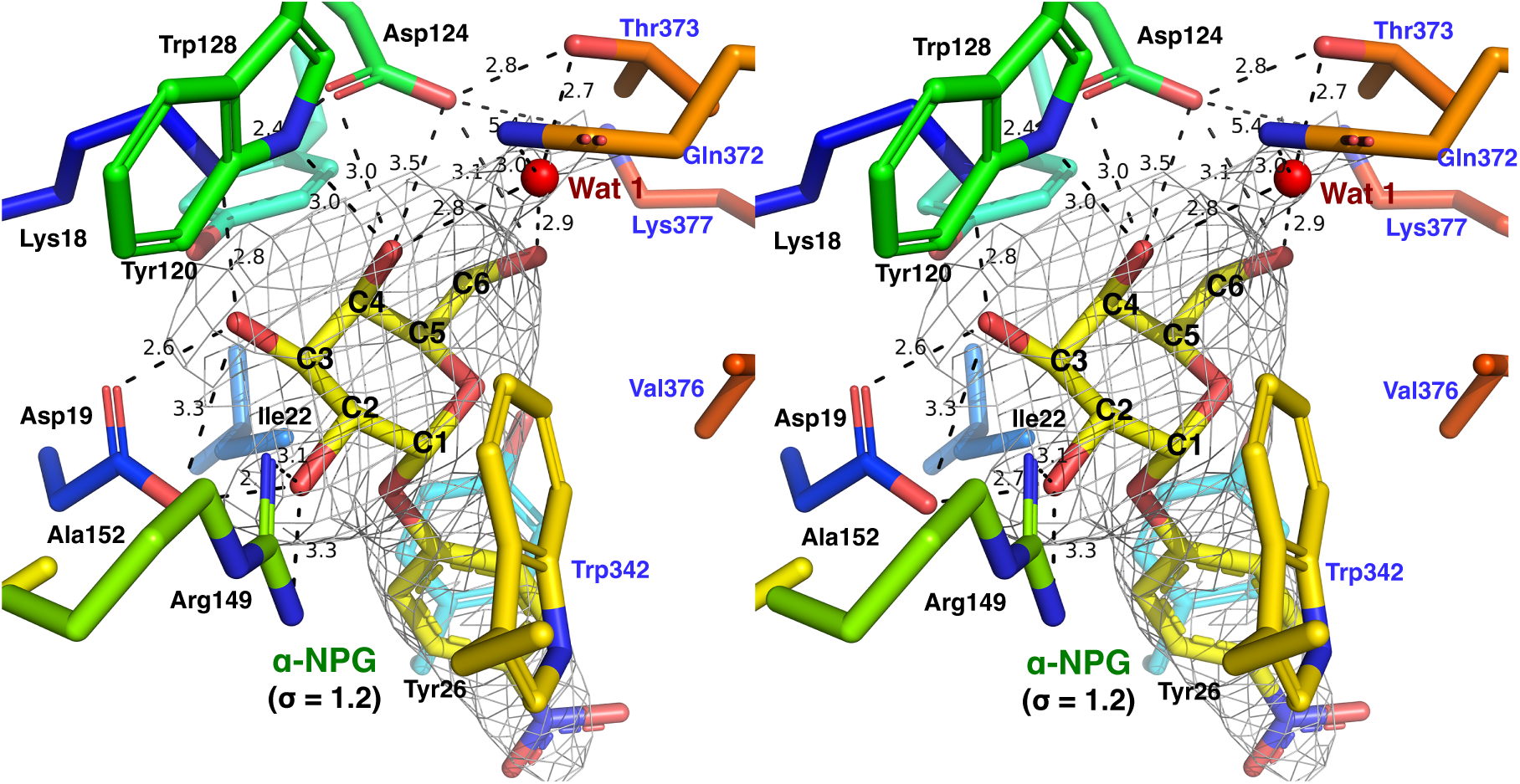
α-NPG binding. The cross-eye stereo view of α-NPG binding. The side chains forming the binding pocket formed from N- and C-terminal domains were shown in stick representation, colored according to corresponding hosting helices in rainbow, and labeled in black and blue, respectively. Isomesh map of the α-NPG (in yellow) and Wat 1 (in red) were contoured at a level of σ =1.2. Dashed lines indicate distances within hydrogen-bonding or salt-bridge interactions (Å).

#### Melibiose binding

The endogenous substrate melibiose is formed from a galactose unit and a glucose unit linked by an α-1,6 galactosyl bond (D-Gal-(α1→6)-D-Glc). The melibiose-bound D59C MelB_St_ structure was refined to a resolution of 3.05 Å (**Fig. 4a, e**), and the density map displayed a clear two-unit blob, fitting well with one molecule of melibiose. Two water molecules, Wat 2 and Wat 3, were modeled; however, no strong peak was observed at the Wat 1 position. Notably, the binding affinity between melibiose and α-NPG differs by a factor of 100; in α-NPG, the hydrophobic phenyl group, positioned at 4 Å distance from the phenyl group of Tyr26, forms a strong stacking interaction. While melibiose has four additional OH groups, the glucosyl ring sits nearly perpendicular to the phenyl ring of Tyr26 **(Fig. S3**) with no strong polar interaction with the transporter.

**Fig. 4.**
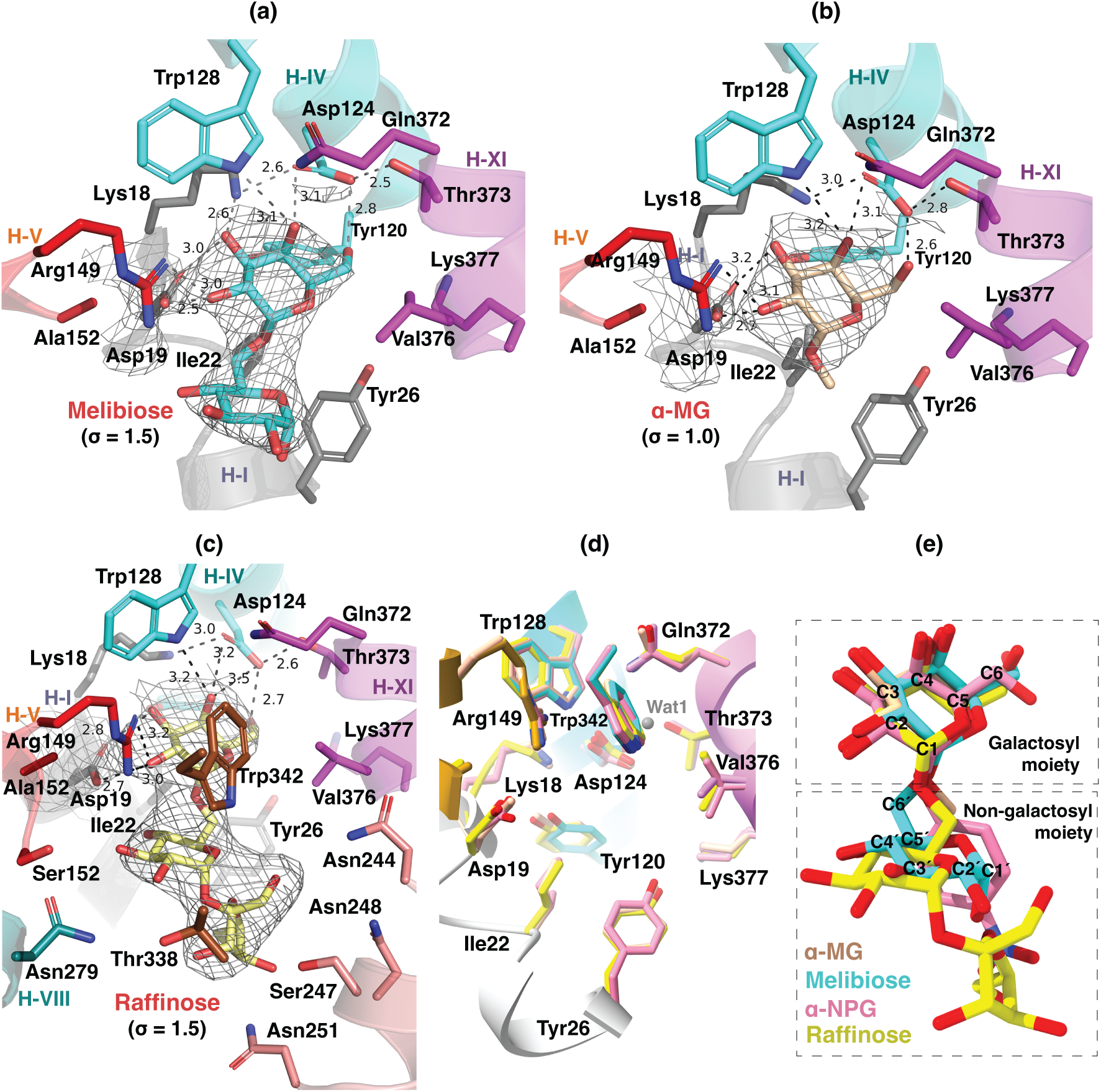
Binding of melibiose, α-MG, and raffinose. All residues within a 5-Å distance to the bound substrates were shown in sticks. Dashed lines, the distances within hydrogen-bonding and salt-bridge interactions (Å). **(a)** Melibiose binding. **(b)** α-MG binding. **(c)** Raffinose binding. **(d)** Residues in the sugar-binding pockets from the alignment of all four structures. **(e)** Substrates from the alignment of all four structures. Carbon positions on the galactosyl (C1-6) and glucosyl moiety (C1′-6′) were labeled on the melibiose molecule. Trp342 was removed for clarity in panels a, b, and d. Isomesh maps for each sugar and Asp19 were contoured at levels of σ = 1.5 for melibiose and raffinose or σ = 1.0 for α-MG.

### Methyl α-D-galactoside (α-MG) binding

α-MG has a methyl substituent at the anomeric C1 position of the galactosyl moiety in an α-linkage. The crystal structure of D59C MelB_St_ complexed with α-MG was refined to a resolution of 3.68 Å, and the density map clearly displayed a one-unit blob in the binding pocket, where one α-MG was modeled similarly to the galactosyl moiety of melibiose (**Fig. 4b**).

#### Raffinose binding

The raffinose-bound D59C MelB_St_ mutant structure was refined to a resolution of 3.40 Å, and the density map displayed a clear three-unit blob in the binding pocket, fitting well with this trisaccharide. As expected, the raffinose-binding pocket is significantly larger than that of melibiose and α**-**NPG **(Fig. 4c)**, involving two additional helices. Consequently, a set of polar side chains on helix VII (Asn244, Ser247, Asn248, and Asn251), helix VIII (Asn279), and helix V (Ser153) were within 4-5 Å distances to either the glucosyl or fructosyl moieties. Interestingly, the polar interactions from raffinose to MelB_St_ are limited to the galactosyl moiety, which are nearly identical to those presented in α-MG, melibiose, or α-NPG, while the Asp19 is at a hydrogen-bonding distance from the OH-4 on the glucosyl moiety.

All four galactosyl moieties align well in the specificity-determinant pocket, regardless of the number of monosaccharide units (**Fig. 4d, e; Fig. S1**), which provided a strong structural support to the previous conclusion that the galactosyl moiety determines the specificity of the primary substrates and the non-galactosyl moiety contributes to the binding affinity.

### Structural dynamics measured by HDX-MS

To determine the structural dynamics of MelB_St_ and the influence of melibiose and/or Na^+^, the differential HDX-MS experiments were employed between MelB_St_ alone and MelB_St_ with melibiose and/or Na^+^ (28, 32, 33) (**Table S2**). MelB_St_ residues 2-470 out of 476 were covered, achieving a coverage of over 87.47% and 86.62% from 149 or 152 overlapping deuterium-labeled peptides (**Tables S2-3; Fig. S4**). There are 59, 63, or 59 non-covered residues for the datasets of the apo vs. melibiose-bound, Na^+^-bound, and melibiose- and Na^+^-bound states, respectively. Most are at the transmembrane helices IV, IX, X, and XI, and some cytoplasmic loops. Labeled peptides cover all periplasmic loops.

#### Six regions with greater deuteriation at the apo state

The peptide coverage-based deuteriation per residue plot and the relative deuterium uptake per peptide were generated from mean values from all three time points of each peptide with six duplicates (**Fig. 5a,b; Table S3**). The data indicate that, aside from the N- and C-terminal tails, which exhibit high HDX, six regions showed greater deuterium uptake in the apo state. As highlighted in red, two were in the N-terminal helices (I and V) and four were in the C-terminal loops (loop_6-7_, loop_8-9_, loop_9-10_, and loop_10-11_), which are the dynamic regions of MelB_St_ in the apo state. Notably, all the ICH1-3 were in the dynamic area. Comparing with helices I and V carrying the sugar-binding residues, helices II and IV—housing both the Na^+^-binding residues (Asp55, Asn58, and Asp59) and part of the sugar-binding residues (Asp124 and Trp128)—exhibited significantly lower deuteriation levels.

**Fig. 5.**
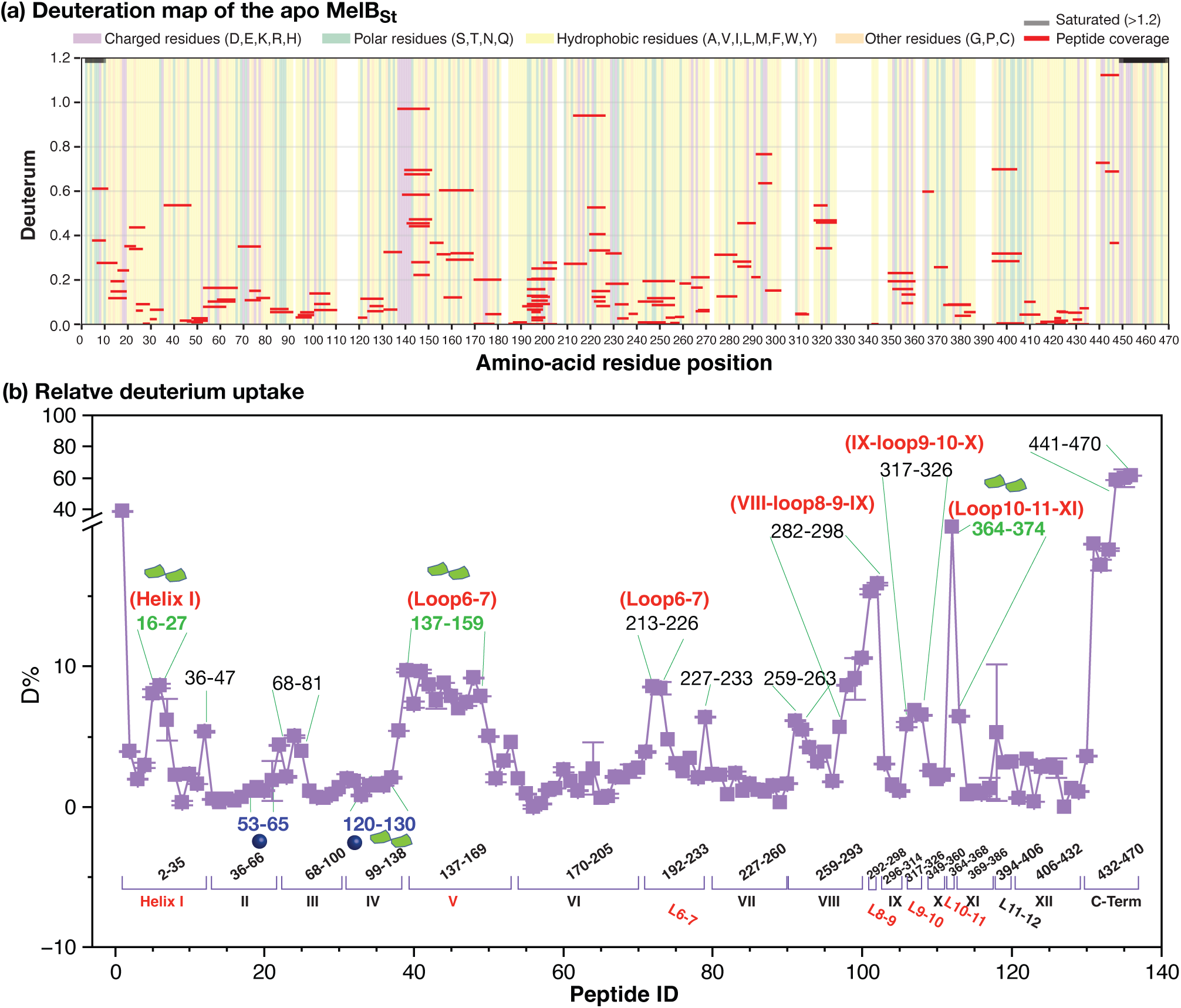
Deuterium uptake of the apo MelB_St_. HDX experiments on WT MelB_St_ were conducted as described in Methods. All values were from the mean of six measurements at the apo state. **(a)** Deuteriation map of the apo MelB_St_. Mean deuteration levels of MelB_St_ peptides at the apo state averaged first across timepoints (30s, 300s, 3000s), then across two replicates, were presented against amino-acid residue sequence. The values at both N- and C-terminal regions were greater than 1.2 showed as black bars. The chemical properties of the peptide-covered residue were indicated by background shading; the white background indicated the non-covered positions. **(b)** Relative deuterium uptake per peptide plot. The corresponding transmembrane helices and a few loops were marked. Notably, due to the nature of overlapping peptides, the indicated amino-acid position is not sequential. Peptides with greater deuterium uptake were labeled. Peptides covering sugar- and cation-binding sites were colored in green and blue, respectively, and the dynamic regions with greater uptakes were colored in red.

#### The effects of substrate(s) on MelB_St_ dynamics

The differential deuterium labeling (ΔD), which was calculated based on deuterium uptake in the absence (apo state) or presence of specific ligand(s) (Holo state), was shown as the residual plot of each peptide at three labeling time points and the sum of all (**Fig. 6a**). The dashed lines indicate the global thresholds calculated for each dataset. A hybrid significance analysis was used to determine the significance: ΔD > the global threshold of each dataset and P < 0.05 (34). More than 50% of residues (237, 264, or 257 positions) exhibited either insignificant ΔD values (D_Mel-Apo_ < |0.186|, ΔD_(Na+)-Apo_ < |0.224|, or ΔD_Na(+)Mel-Apo_ < |0.175|), or P > 0.05, respectively (**Table S2**). Melibiose binding induced a wide range of effects on HDX, including a few protections (less deuterium uptake in the Holo state) and more deprotections (greater deuterium uptake in the Holo state). In contrast, Na^+^ alone or with melibiose primarily caused deprotections. The results supported the previous conclusion determined by the thermal denaturation study detected by circular dichroism spectroscopy (30); i.e., the melibiose-, or Na^+^-bound MelB_St_ was more stable than the apo state, and when MelB_St_ bound with both, it was the best. All deprotected peptides with significance were labeled individually (**Fig. 6a**). A peptide with at least one significant ΔD (meeting both criteria) from any time points was mapped onto the melibiose-bound structure (**Fig. 6b-d; Fig. S4b**).

**Fig. 6.**
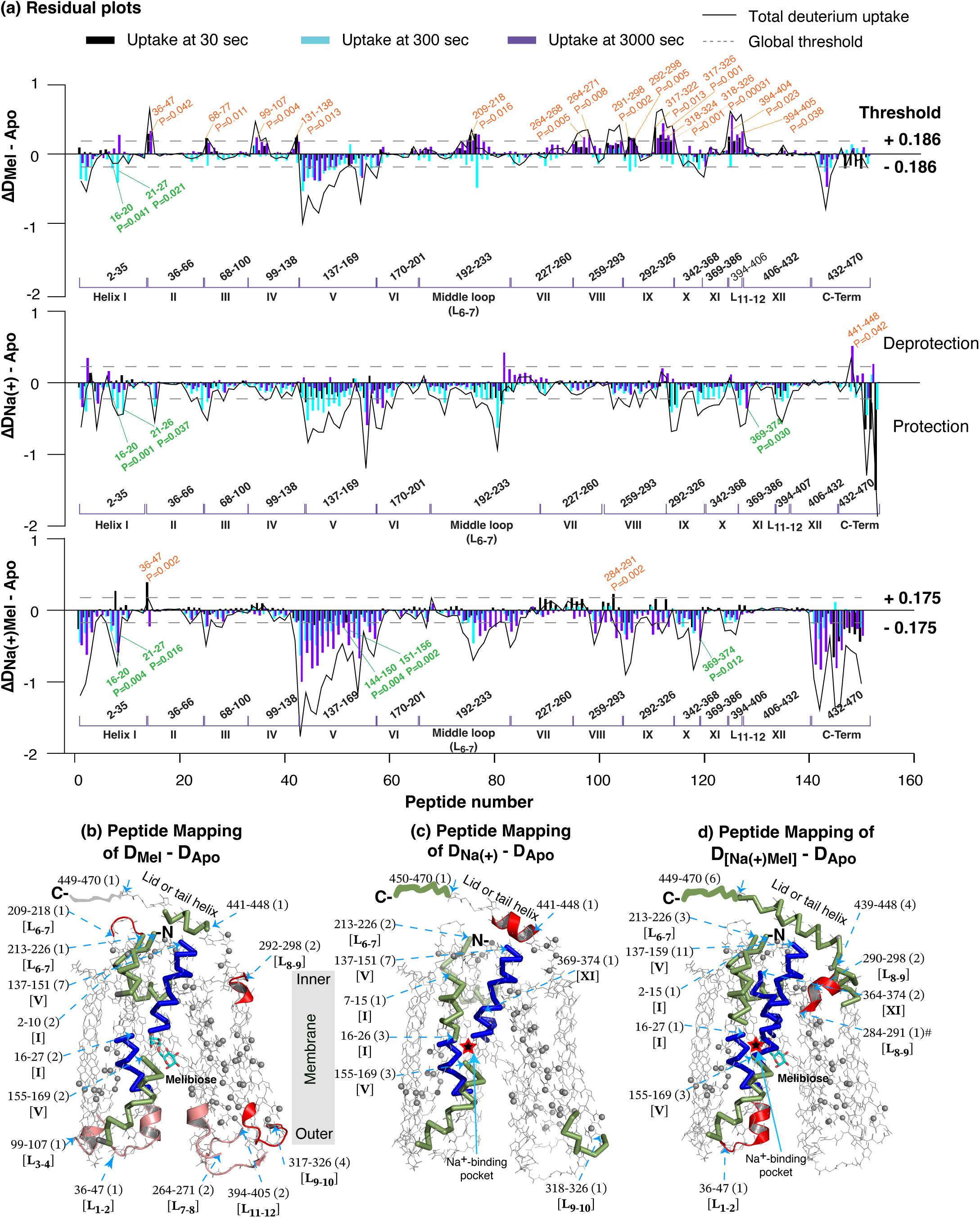
Residual plots and structural mapping. HDX experiments on WT MelB_St_ in the apo or holo states (with melibiose, Na^+^, or Na^+^ plus melibiose) were conducted as described in Methods. **(a)** Residual plots (D_Holo-Apo_). Differential deuterium uptakes (ΔD) of each time point and the total uptake calculated from paired conditions for position 2-470 were plotted against the peptide number. Black, cyan, and purple bars, the deuterium uptake at 30, 300, and 3000 sec, respectively; dark gray curve, total uptake from all three time points. Deprotection, ΔD_Holo–Apo_ > 0; protection, ΔD_Holo–Apo_ < 0. Each sample was analyzed in triplicate. Dashed lines indicate the levels of the global threshold values calculated from each dataset, as labeled. The protein residue positions corresponding to the overlapping peptides were marked, and the covered transmembrane helices were labeled in Roman numerals. All deprotected peptides with statistical insignificance (ΔD >threshold and P<0.05) in each dataset were labeled. **(b-d)** Peptide mapping on the crystal structure of the melibiose-bound inward-facing conformation for D_Mel–Apo_, D_Na(+)–Apo_, and D_Na(+)Mel–Apo_, respectively. The red star symbol indicated the location of the Na^+^-binding pocket, the drawing lines represented the disordered MelB_St_ C-terminal tail, and a gray bar showed the membrane region of MelB_St_. The peptides of ΔD values with statistical significance (ΔD > |threshold| and P < 0.05) at any timepoint are highlighted either in ribbon representation for protection (colored in blue for peptides covering sugar-binding pocket and in green for all other regions) or in cartoon representation for deprotection (colored pink for data from 3000 sec and red for data from 30 sec). Non-covered residues from each dataset are shown as gray spheres at the Cα position and listed in **Table S3**, and peptides with statistically insignificant differences (either ΔD < |threshold| or P>0.05) are illustrated in backbone representation in gray. Peptide positions are marked by their starting residue; the number of overlapping peptides is indicated in round brackets, and the location is shown in square brackets.

Consensus effects were observed in all three labeling conditions (**Fig. 6a-d**), and peptides with significant effects were clustered in several regions, which largely overlapped with the dynamic regions obtained at the apo state (**Fig. 5**). Four out of six dynamic regions showed protection. Nearly full-length helices I and V, and regions 213-226 covering ICH1, showed protection by melibiose and/or Na^+^. The peptides covering 364-374 at the loop_10-11_ and the starting part of helix XI, which contains the sugar-binding residues Gln372 and Thr373, were protected by Na^+^ or Na^+^ with melibiose. Another two dynamic regions exhibited diverse effects. The cytoplasmic positions 292-298 covering the loop_8-9_ and the starting helix IX, containing a conformation-important residue Arg295, were deprotected by melibiose but protected by melibiose and Na^+^. The neighboring peptide 284-291, corresponding to helix VIII and loop_8-9_ covering ICH2 at the cytoplasmic gating area, was also deprotected at the 30-sec time point but protected at the 3000-sec time point by melibiose and Na^+^ (**Figs. 6d**). The positions 317-326 at helix IX-loop_9-10_-helix X showed deprotections by melibiose but were protected by Na^+^. Five out of six loops, except for loop_5-6_ (**Fig. 6a-b, pink and red**), were deprotected by melibiose. In addition, the lid ICH3 was protected by melibiose or melibiose with Na^+^, with a single result of deprotection by Na^+^. The following sections will focus on the sugar- and cation-binding pockets as well as the structural elements critical for conformational transitions. All peptides with ΔD values greater than the threshold value were selected, and their deuterium uptake time course plots were presented in the Supplemental Figure S5.

#### A flexible sugar-binding pocket with a rigid cation-binding pocket

This HDX study covered most positions for the binding pockets for sugar and Na^+^ (**Table S4)**. Ten representative peptides were highlighted by uptake time course plots with structural mapping (**Fig. 7**). Three peptides 53-62 (helix II) and 120-123 & 121-130 (helix IV), which cover all Na^+^-binding residues (Asp55, Asn58, Asp59, and Thr121), exhibited lower deuteriation levels at the apo state (**Figs. 5, 7, black open squares**) with no significant effect by melibiose and/or Na^+^ (**Fig. 7, blue filled squares)**. Others in helices II and IV also exhibited similar behavior (**Fig. 5; Table S3**), indicating that both helices, including the Na^+^-binding residues, are conformationally rigid.

**Fig. 7.**
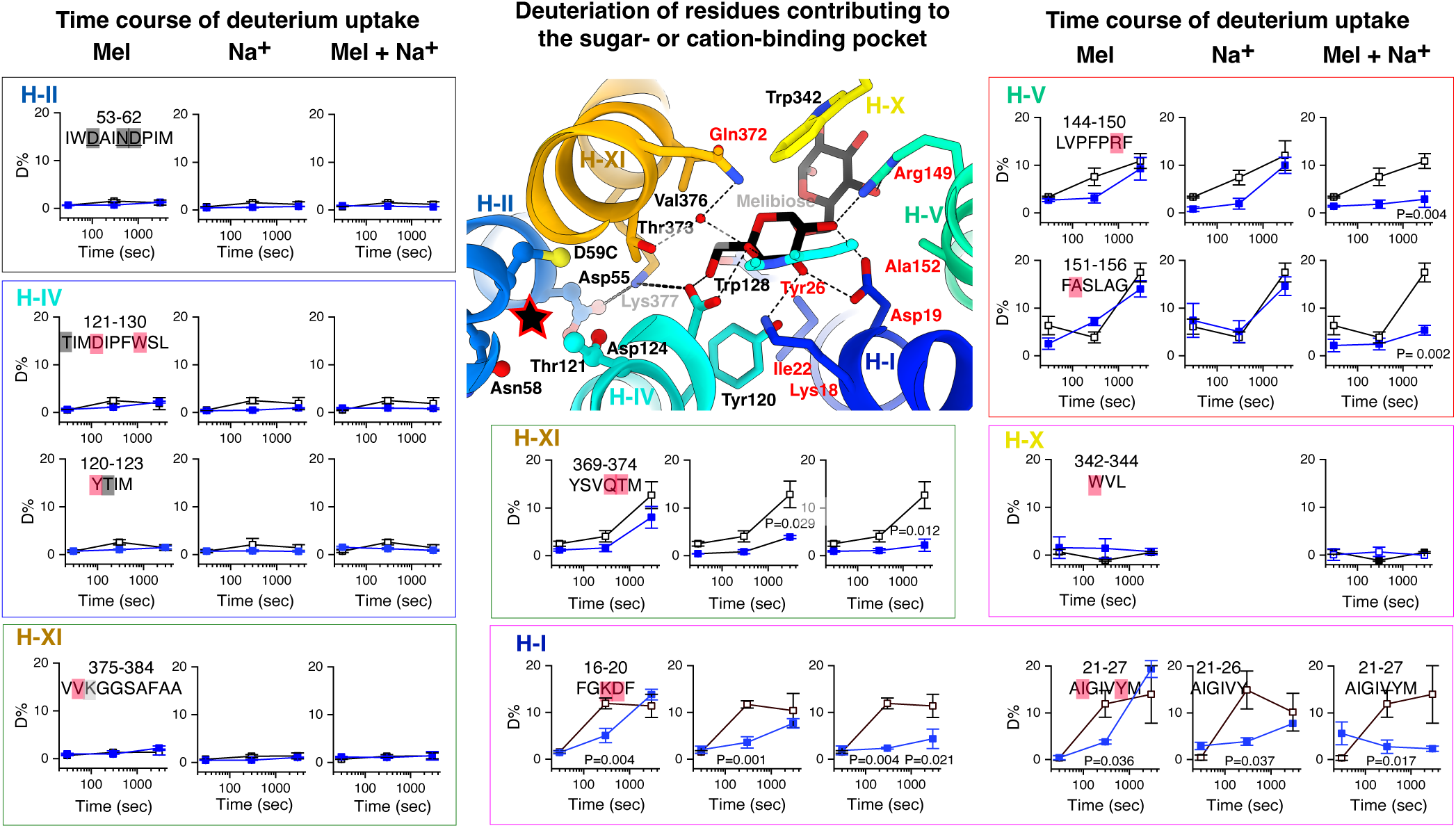
Allosteric effects on substrate-binding sites. Deuterium uptake time course of representative peptides covering the sugar- and cation-binding pockets. The percentage of deuterium uptake measured in the absence (empty black square) or presence of melibiose (Mel), Na^+^, or Mel & Na^+^ (filled blue square) was plotted against labeling times of 0, 30, 300, and 3000 sec. The peptide sequences were shown with residues at the cation- or sugar-binding pocket highlighted in black or red, respectively. Lys377 between the two binding pockets was highlighted in gray. P values are provided for each time point where the ΔD value exceeds the threshold. On the melibiose-bound structure, residues participating in the sugar binding and cation binding were shown in stick. Red start, the cation-binding pocket; red text labels: residues showing greater HDX with significant substrate effects; black text labels: residues showing poor HDX with no significant substrate effects.

As described, helices I and V are the major flexible transmembrane helices (**Fig. 5**), which cover the six sugar-binding residues (Lys18, Asp20, Ile22, Tyr26, Arg149, and Ala152). Peptides 16-20 and 21-27 (helix I) were significantly protected by melibiose, Na^+^ that binds remotely, or both (**Fig. 7)**. Arg149 (helix V) was well covered by several overlapping peptides that consistently demonstrated greater protections by melibiose and/or Na^+^ (**Fig. 6-7**). The shorter peptide 144-150 (helix V) showed protection by melibiose or Na^+^ alone, which is statistically significant; however, the magnitude of the change is subtle. In the presence of both, the protection became significant. The Ala152-carrying peptide showed significant protection by melibiose and Na^+^, and peptide 368-273, which covers the sugar-binding residues Gln372 and Thr373 (helix XI), also exhibited significant protection by Na^+^ alone or in combination with melibiose. Peptides carrying other sugar-binding residues Asp124, Try128, and Trp342 (helix IV) and Val376/Lys377 (XI) showed poor deuteriation and no significant effects by either melibiose, Na^+,^ or both. Overall, the results indicated that the sugar-binding residues in proximity to the cation-binding pocket are rigid with no significant effect by either substrate; in contrary, peptides carrying the sugar-binding residues far from the cation-binding site are dynamic, and their flexibility were significantly inhibited by sugar binding itself, by Na^+^ alone, especially by the binding of both, which support that the melibiose affinity is correlated with the conformational flexibility and Na^+^ can increase the sugar binding.

#### Dynamics of the cytoplasmic gating salt-bridge network and modulations by substrates

The cytoplasmic gating salt-bridge network between the two domains, involving nine charged residues (**Fig. 8a, c**), was most covered. Residues Lys138, Arg141, and Glu142 at loop_4-5_-helix V are three critical positions from the N-terminal domain. Overlapping peptides that covered this region consistently showed greater deuteration levels at the apo state and greater protections under all three conditions, as represented by the peptide 137-150 **(Figs. 5-7**). Notably, the sugar-binding residue Arg149 at the gate area of helix V is within this most dynamic area of the transmembrane domain of MelB_St_ and the same dynamics group. Another gating residue, Arg295 (helix IX), which forms multiple interactions in this network, including two water molecules (**Fig. S2**), was covered by peptide 292-298, which showed a higher deuteration level and was deprotected by melibiose but protected by melibiose and Na^+^. Peptide 364-368 at Loop_10-11_ and the connected region of helix XI, which contains Glu365 at the other side of this salt-bridge network, also showed high deuterium uptake and protection by melibiose with Na^+^. On the contrary, Asp341, Asp354, and Glu357 at the rigid helix X, which are located in the center of this salt-bridge network, showed poor deuteration under all conditions. The results showed that gating residues surrounding the rigid helix X in this broadly spanned cytoplasmic salt-bridge network are highly dynamic, and sensitive to the binding of melibiose, Na^+^, and melibiose with Na^+^.

**Fig. 8.**
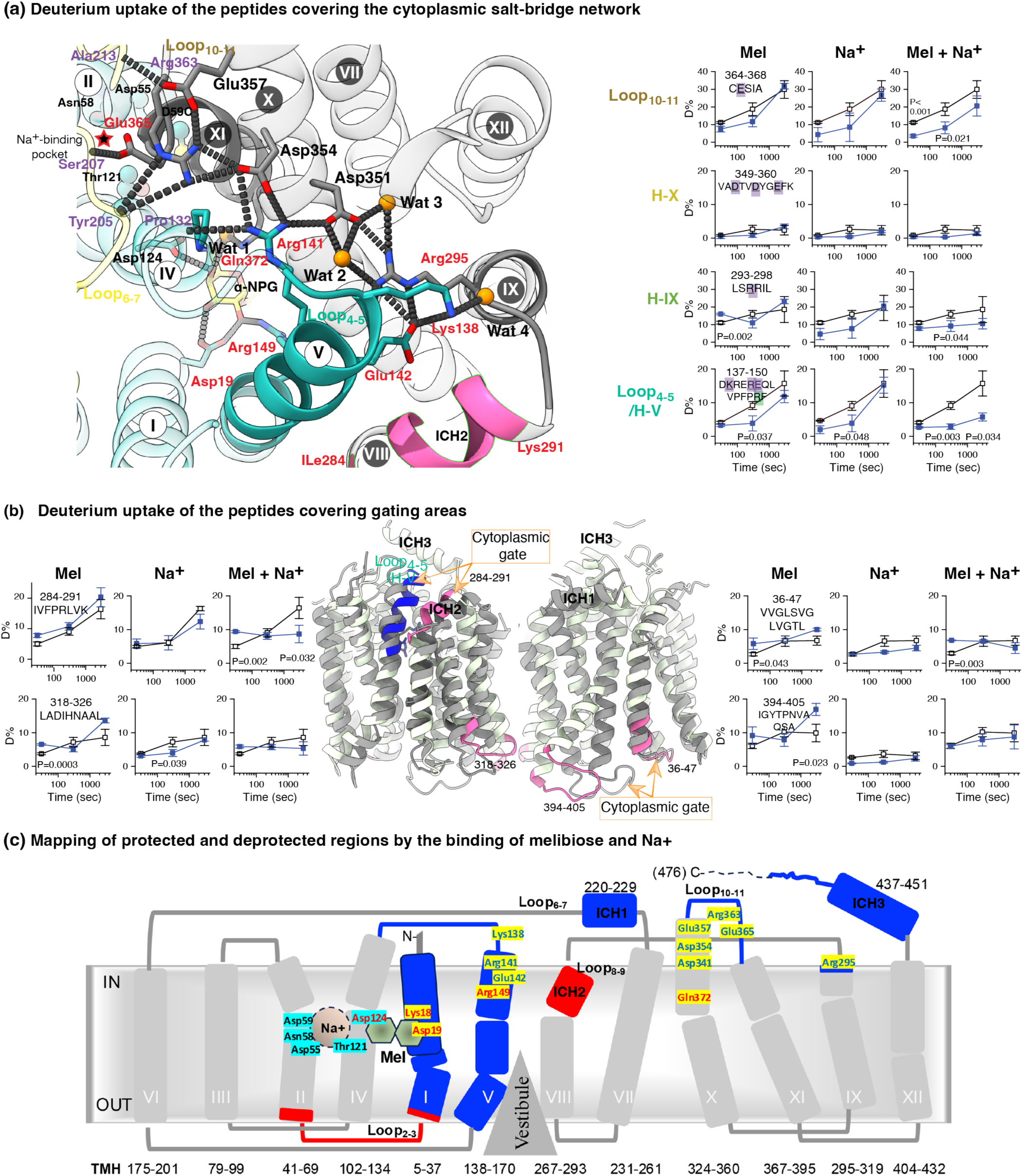
Allosteric effects on gating salt-bridge network. **(a)** Deuterium uptake time course of representative peptides covering the cytoplasmic gating salt-bridge network. The percentage of deuterium uptake measured in the absence (empty black square) or presence of melibiose (Mel), Na^+^, or Mel & Na^+^ (filled blue square) was plotted against labeling times of 0, 30, 300, and 3000 sec. P values are provided for each time point where the ΔD value exceeds the threshold. The peptide sequences were shown with residues at the salt-bridge network or sugar-binding pocket highlighted in purple or red, respectively. On the α-NPG bound structure, the gating salt-bridge network (Lys138, Arg141, and Glu142 (Loop_4-5_/IV), Arg295 (IX), Asp351, Asp354, and Glu357 (X), and Arg363 and Asp365 (Loop_10-12_) and their polar contacts with the backbone of other positions were shown in dashed lines. The N- and C-terminal domains were colored in light cyan and gray, respectively. The residues at the cation binding pocket were shown in ball and stick, as also indicated by the red star. Wat, water. Three sugar-binding residues, Asp19, Arg149, and Gln272, were shown in stick. Arg363 is in the list of uncovered positions. The region covering both the cytoplasmic gating salt-bridge network and sugar-binding residue Arg149 was highlighted in solid color, and the peptide 284-291 at helix III-ICH2 was also highlighted in pink. **(b)** Deprotection at loops. Deuterium uptake time course of four peptides at loops, mainly at the gating area, was presented and also mapped on the outward-facing structure, which was overlayed with the inward-facing structure [PDB 8T60]. The cytoplasmic and periplasmic gates were indicated. The peptides with deprotection by substrate were colored in pink. **(c)** HDX and ligand effects mapping on an outward-facing topology model of MelB_St_. Melibiose-and Na^+^-binding residues were labeled in red or black, respectively. Residues showing ligand-induced protection and deprotection were colored in blue or red, respectively.

As shown by the aligned outward- and inward-facing conformations and the membrane topology (**Fig. 8b-c**), two gating regions at ICH2 and helix I-loop_1-2_-helix II, covered by C-terminal peptide 284-291 and N-terminal peptide 36-47 at the cytoplasmic or periplasmic side, respectively, were deprotected by melibiose or together with Na^+^. In addition, another gating region at loop_11-12_ covered by the C-terminal peptide 394-405 and the peptide 318-326 at helix XI-loop_9-10_-helix X was also deprotected by melibiose. Most deprotected areas are at the periplasmic loops.

### Molecular dynamics (MD) simulations

Three systems, including the apo, melibiose-bound, and melibiose- and Na^+^-bound states of a generated WT MelB_St_, were constructed by embedding in a POPE:POPG (7:2) lipid bilayer. For each system setup, five independent replicas of MD simulations were carried out, each for ∼400 ns, and the total sampling time for each system was ∼2 μs.

#### Water 1 occupancy

To analyze the Wat 1 modeled in the higher resolution structure with α-NPG bound, a distance-based criterion for defining the average water occupancy in the sugar-binding site across all trajectories of each system as described in Methods. Results showed that Wat-1 exhibited nearly full occupancy when melibiose was present, regardless of whether Na^+^ was bound at the cation-binding site (**Table S6**), which supported the crystal structure observation.

#### Side-chain flexibility

The side-chain heavy-atom root-mean-square fluctuation (RMSF) over five replicas of trajectories was analyzed for the apo and MelB_St_ with melibiose and Na^+^ bound states (**Fig. 9**). The results showed that the side chains in the cation-binding site and nearby sugar-binding positions educed fluctuations upon the binding of substrates. The residues Asp19, Tyr26, Arg149, and Gln372 at helices I, V, or XI, respectively, were relatively flexible, and their conformational freedoms were significantly reduced by the binding of melibiose and Na^+^. The data about the sidechain flexibility are consistent with the peptide-based HDX results.

**Fig. 9.**
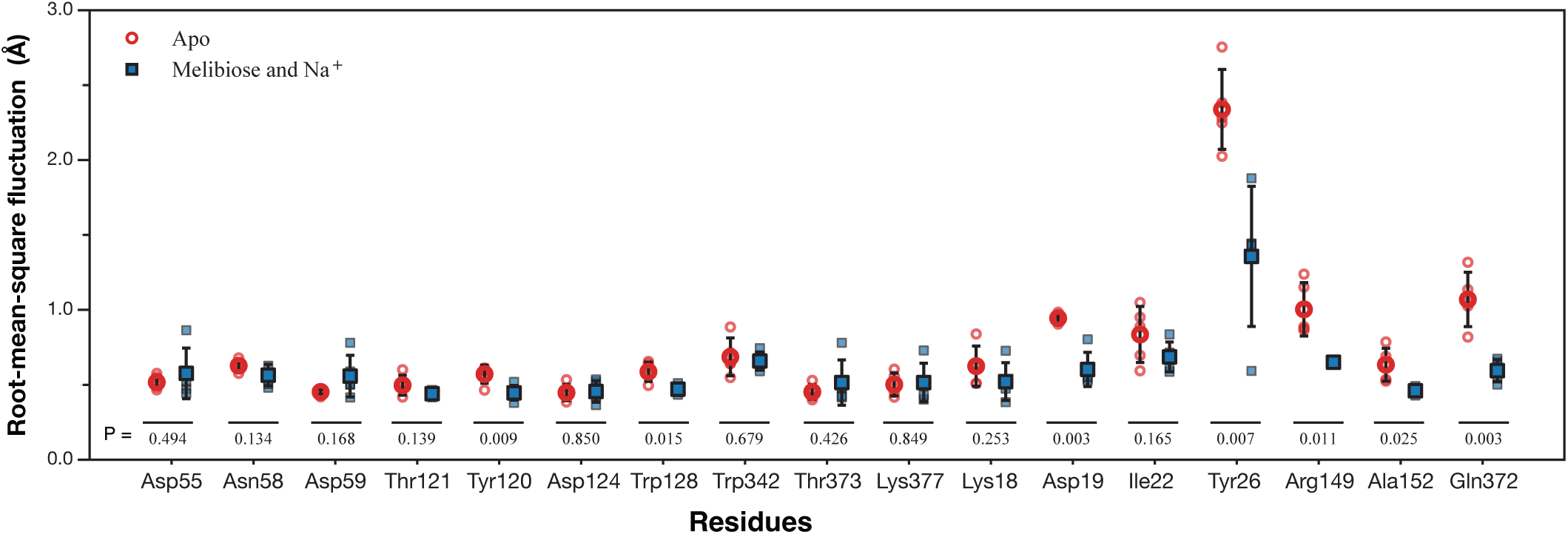
Side-chain flexibility analyzed from MD trajectories. For each residue, the side-chain RMSF values in each of the five replicas for the apo and melibiose- and Na^+^-bound states were plotted. The mean of the RMSF values in each state is represented as red circle (apo state) or blue square (melibiose- and Na^+^-bound state). For each residue, unpaired t test of the RMSF mean values between the apo and the melibiose- and Na^+^-bound states were performed and the p-values were presented.

## Discussion

### The binding recognition and affinity of MelB_St_ have been further refined

Dehydration of sugar molecules is expected for binding; however, full or partial dehydration in MelB is unknown. The 2.60 Å resolution α-NPG-bound structure showed a partially dehydrated sugar at the binding site. The bound water connected the OH-4 and OH-6 on the galactopyranosyl ring with Thr373 and Gln372 at helix XI, and it was also surrounded by Asp124 at helix IV. Previous Cys-scanning mutagenesis has shown that the T373C mutant retained most activities. Still, a Cys residue on the Gln372 position significantly decreased the transport initial rate, accumulation, and melibiose fermentation, with little effect on the protein expression (35), which supported the role of Wat-1 in the binding. Notably, the orientation of OH-4 is crucial for distinguishing between galactose and glucose. Previously, the OH-4 has been shown at hydrogen-bonding distances from the carboxyl group of Asp124 and the indole group of Trp128 at Helix IV (20, 22), and the identification of Wat-1 in this study added the polar residues Gln372 and Thr373 on helix XI to stabilize the critical OH-4 (**Fig. 3**). Thus, these interactions define the specificity of galactosyl molecules and Wat-1 is likely part of sugar binding. Notably, the OH-4 and OH-6 and Wat 1 are in close proximity to the Na^+^-binding residue Thr121 and the cation-binding important residue Lys377, implying that the Wat-1 is between the two specificity-determining pockets. As shown previously, the OH-3 and OH-2 at the opposite edge of the galactopyranosyl ring form multiple hydrogen-bonding interactions with the charged residues Asp19 and Arg149 at helices I and V, which can be assigned to play a crucial role in stabilizing the recognition of OH-4 and enhancing the binding affinity.

#### Dynamics of MelB_St_ at different regions

HDX-MS is a powerful technique that simultaneously discloses the dynamic information in various areas of a protein. The qualitative method overcomes the drawbacks of most site-specific labeling-based techniques; however, it only provides the conformational dynamic information on the equilibrium of all ensembles. In the current study, we determined the deuterium uptake rates of the full-length MelB_St_ in the absence or presence of melibiose and/or Na^+^, identified major dynamic regions (**Fig. 5**), as well as substrate-induced effects on the substrate-binding sites and structural elements critical for conformational transition, including the cytoplasmic salt-bridge network and the gating areas (**Figs. 6-8**).

Our studies of HDX compensated with molecular dynamics simulations on side-chain fluctuations showed that the Na^+^-binding pocket and the sugar-binding residues near the Na^+^-binding site at helix IV are conformationally rigid since they exhibited low deuteration level for both unbound and bound states (**Figs. 5-7, 9; Tables S3-4**), which are consistent to the previous HDX-MS results of conformation transition(21). Those rigidities are likely derived from their hosting helices (II and IV). In addition, the Na^+^-binding affinity, different from sugar binding, exhibited little difference between the inward- and outward-facing states (21). The conformational freedom of the cation-binding site and most sugar recognition positions was likely restricted in most states to facilitate sugar binding.

The sugar-binding residues, located away from the Na^+^-binding site, are dynamic, exhibiting higher deuterium levels and side-chain fluctuations. Most of those positions are within the six dynamic regions identified from the apo state, including Lys18, Asp19, Ile22, Tyr26, Arg149, Ala152, Gln272, and Thr373 at helices I, V, and XI. Those regions were protected to varying extents by melibiose or Na^+^, and the protection was significantly greater when bound to both substrates (**Figs. 5-7**). Even their conformational flexibility was restrained by melibiose and Na^+^, but the deuterium uptakes at those regions were still greater than those of peptides covering the cation-binding site (**Fig. 7**). Notably, substrate-induced protection was also detected in other areas of the hosting helices I and V, as further stressed below.

#### Arg149, a sugar-binding residue in the major dynamic gating area

Abundant peptides covering Arg149 consistently showed protection by melibiose or Na^+^, especially when both were present (**Figs. 5-8**). Structurally, Arg149 is near to a stretch of charged residues as shown by this peptide 137-DK^137^RER^141^E^142^QLVPFP**<u>R</u>**F-150 (**Figs. 5-7**), where the Lys138, Arg141, and Glu142 have been determined to play critical roles in the functionally crucial cytoplasmic salt-bridge network (**Fig. 8**). The dynamics of Arg149 are likely modulated by this charged network and belongs to a same dynamics group, which provided the structural basis for coupling between the sugar-binding affinity and protein dynamics. At the inward-open structure, this network is deformed, Arg149, as part of this gate, exhibits a large displacement due to an otherwise stereo collision with helix X at the outward-facing state (**Fig. S6**). The sugar-binding affinity at this inward-facing state was reduced by ∼30-fold due to the broken binding pocket and the displacement of Arg149 and Gln372. Functionally, the single-site R149C mutant inhibited but did not eliminate the melibiose transport (18), and single-Cys149 (35) also fermented melibiose with intact cells. The neighboring P148L mutant decreased transport *V*_max_ with a better *K*_m_ (36). All data showed that the dynamics of this charged network and sugar-binding affinity are interconnected.

This cytoplasmic gating salt-bridge network has a rigid center dictated by helix XI and dynamic surroundings from helices V, IX, XI, loops 4-5, 8-9, 6-7, 10-11, and those dynamic positions were significantly inhibited by melibiose, Na^+^, especially the binding of both. Notably, Nb725 binding to the inward-facing conformation also inhibited the dynamics of the similar regions as shown by peptides 137–150, 284–289, and 364–368 (21). The similar HDX profiles from all testing conditions suggested the protein motions followed a similar pathway of the conformational transition. Thus, the dynamics of Arg149 are linked to motion and conformational transitions, as well as sugar-binding affinity. Connectively, it is postulated that Na^+^ binding between helices II and IV stabilizes the dynamics of the gating salt-bridge network and the flexibility of Arg149 and Asp19, which are located more than 10 Å away, thus switching the primary substrate-binding site to the higher-affinity state allosterically.

The binding of sugar further stabilizes this network and promotes Na^+^ binding by preventing Na^+^ from leaving the cation-binding pocket, thus increasing Na^+^-binding affinity (18, 26). Therefore, the cooperative binding of sugar and Na^+^ is part of the substrate-induced conformational allostery, and all functions together.

There are a few regions with substrate-induced deprotections. Two gating areas were deprotected by melibiose and Na^+^, including the cytoplasmic ICH2 and Loop_1-2_, which suggested the structural arrangements. Notably, ICH2 is linked with Arg295 at one end of the salt-bridge network, and the dynamics of this area could influence the stability of the gating network, which might trigger the separation and conformational transition to the inward-open state. Thus, the increased dynamics of loop_1-2_ and Loop_8-9_ at both gating areas by melibiose, especially with Na^+^, can be interpreted as the tendency to form a transition-competent conformation with closing at the periplasmic gate and opening at the cytoplasmic gate. A region at loop_6-7_ responding to the ligand binding identified by the atomic force microscopy was not observed in this HDX study (37).

The symporter H^+^-coupled lactose permease LacY and xylose permease XylE also show protonation dependence of sugar-binding affinity(32, 38, 39), with separate cation- and sugar-binding sites (5, 40–43). Both transporters favored inward-facing conformation and substrate binding induced outward-facing conformation, as indicated by the extensive HDX analysis with XylE (32, 44) and a bunch of biophysical measurements of LacY (45). Notably, several dynamic regions in MelB_St_ were also identified in XylE; both proteins showed binding-induced structural changes in their loop_1-2_ and loop_8-9_ (44). In MelB_St_, while no quantified information on the ensembles, all experimental results and molecular simulations on the minimum free-energy landscape for sugar translocation suggest that the apo MelB favors the outward-facing conformation, and the outward-facing conformation is further preferred when bound with melibiose and Na^+^ (20, 21, 26).

### Summary

Our studies on structure, HDX, and MD simulations allow us to conclude that the sugar-binding affinity of MelB_St_ is coordinated with protein structural motions and conformational transition between inward- and outward-facing states. Na^+^ binding restrains the dynamics of remote sugar-binding residues via stablizing the dynamic cytoplasmic salt-bridge network, thereby increasing sugar-binding affinity allosterically. The conclusion provides insightful knowledge to understand cooperative binding and symport mechanisms.

## Experimental Procedures

### Reagents

Melibiose was purchased from *Acros Organics* (*ThermoFisher Scientific*). [^3^H]Raffinose and label-free raffinose were purchased from American Radiolabeled Chemicals (ARC), Inc and Research Products International (RPI) Corp., respectively. Raffinose (RPI Chemicals), α-methyl galactoside (α-MG) and *p*-Nitrophenyl α-D-galactoside (α-NPG) were purchased from Sigma-Aldrich. Detergents undecyl-β-D-maltopyranoside (UDM) and dodecyl-β-D-maltopyranoside (DDM) were purchased from *Anatrace*. *E. coli* lipids (Extract Polar, 100600) were purchased from *Avanti Polar Lipids, Inc*. All other materials were reagent grade and obtained from commercial sources.

### Strains and plasmids

*E. coli* DW2 cells (*melA*^+^*B^-^, lacZ^-^Y^-^*) (46) were used for protein expression and functional studies. The expression plasmids pK95/ΔAH/WT MelB_St_/CHis_10_(15) and pK95/ΔAH/D59CMelB_St_/CHis_10_ (18) were used for constitutive expression.

### MelB_St_ protein expression and purification

Cell growth for the large-scale production of WT MelB_St_ or D59C MelB_St_ was carried out in *E. coli* DW2 cells (18, 47). Briefly, MelB_St_ purification by cobalt-affinity chromatography (Talon Superflow Metal Affinity Resin, Takara) after extraction by 1.5% UDM. MelB_St_ protein was eluted with 250 mM imidazole in a buffer containing 50 mM NaPi, pH 7.5, 200 mM NaCl, 0.035% UDM, and 10% glycerol, and further dialyzed to change the buffer conditions accordingly.

### Protein concentration assay

The Micro BCA Protein Assay (Pierce Biotechnology, Inc.) was used to determine the protein concentration.

### α-NPG transport

MelB_St_-mediated α-NPG downhill transport was detected by the release of the intracellular α-NPG hydrolytic product, *p*-nitrophenol, from *E. coli* DW2 cells expressing high-turnover number α-galactosidase and recombinantly expressed MelB_St_ (Ottina, 1980). *E. coli* DW2 cells carrying the constitutive expression plasmid for the WT MelB_St_ or D59C MelB_St_ uniporter mutant in LB media containing 100 mg/L ampicillin were grown in a 37 °C shaker as described (15). The overnight cultures were diluted 5-fold into fresh LB broth and 100 mg/L ampicillin, and then shaken at 30 °C for 4 h. The expression of α-galactosidase was induced by adding 5 mM melibiose for 1 hour before harvesting the cells for the transport assay. The melibiose-induced cells were washed with 100 mM KP_i_, pH 7.5, three times to remove the remaining melibiose and Na^+^, and adjusted to *A*_420_ of 10 (∼0.7 mg proteins/ml) in the assay solution of 100 mM KP_i_, pH 7.5, 10 mM MgSO_4_, and 1 mM DTT in the absence or presence of 20 mM NaCl or LiCl. The cells under each condition were equilibrated at a 30 °C incubator for 10 minutes before the α-NPG transport assay, which was initiated by mixing 0.5 mM label-free α-NPG and incubating for 60 min. A 100-µL cell aliquot at the given time point of 0, 1, 5, 10, 20, 30, 60 min was quenched with 900-µL 0.33 M Na_2_CO_3_, followed by a centrifugation at 10,000 g for 5 min to collect the clarified supernatant for absorbance measurements. The extracellular *p*-nitrophenol released from the intracellular hydrolytic product of the translocated α-NPG, which was supplied in the extracellular environment, was measured at 405 nm by a UV spectrometer. The concentration of *p*-nitrophenol is estimated based on its molar extinction coefficient of 18000 M^-1^cm^-1^.

### [^3^H]Raffinose transport

The raffinose active transport was carried out by [^3^H]raffinose uptake with DW2 cells expressing MelB_St_ without the induction of α-galactosidase as described for the melibiose transport assay(15). Briefly, the cells expressing the WT MelB_St_ or D59C MelB_St_ uniporter mutant, or without a plasmid, were mixed with 2-µL of 25 mM [^3^H]raffinose (specific activity 10 mCi/mmol) to 50 µL of cells (final raffinose concentration at 1 mM in the absence or presence of 50 mM NaCl or LiCl, and the transport reaction was quenched at the given time points and followed by a fast filtration.

### Isothermal titration calorimetry

All ITC ligand-binding assays were performed with the TA Instruments (Nano-ITC device) as described (19), which yields the exothermic binding as a positive peak. The MelB_St_ was dialyzed overnight with assay buffer containing 20 mM Tris-HCl (pH 7.5), 100 mM NaCl, 10% glycerol, and 0.035% UDM. The ligands are prepared by dissolving in the same batch of dialysis buffer for buffer matching. In a typical experiment, the titrand (MelB_St_) placed in the ITC Sample Cell was titrated with the specified titrant raffinose or α-methyl galactoside (placed in the Syringe) in the assay buffer by an incremental injection of 2 or 2.5-μL aliquots at an interval of 250 or 300 sec at a constant stirring rate of 250 rpm (nano-ITC). MelB_St_ protein samples were buffer-matched to the assay buffer by dialysis. The normalized heat changes were subtracted from the heat of dilution elicited by the last few injections, where no further binding occurred, and the corrected heat changes were plotted against the mole ratio of the titrant to the titrand. The values for the binding association constant (*K*_a_) were obtained by fitting the data using the one-site independent-binding model included in the NanoAnalyze software (version 3.7.5). The dissociation constant (*K*_d_) = 1/*K*_a_.

### Crystallization, native diffraction data collection, and processing

The D59C MelB_St_ dialyzed overnight against the sugar-free dialysis buffer (20 mM Tris-HCl, pH 7.5, 100 mM NaCl, 0.035% UDM, and 10% glycerol), concentrated with Vivaspin column at 50 kDa cutoff, and stored at - 80 °C. A phospholipid stock solution of 20 mM was prepared by dissolving the *E. coli* Extract Polar (Avanti, 100600) with a dialysis buffer containing 0.01% DDM. The protein sample was diluted to a final concentration of 10 mg/ml with the same sugar-free dialysis buffer, supplemented with phospholipids at 3.6 mM and 30 mM of melibiose or α-MG, 40 mM raffinose, or 6 mM of α-NPG in DMSO solution. Crystallization trials were conducted using the hanging-drop vapor-diffusion method at 23°C by mixing 2 μL of protein with 2 μL of reservoir solution. Crystals from D59C MelBSt protein with the melibiose, α-MG, or α-NPG appeared against a reservoir consisting of 100 mM Tris-HCl, pH 8.5, 100 mM NaCl_2_, 50 mM CaCl_2_, and 32-35% PEG 400. For the raffinose-containing sample, the crystals were collected from 100 mM Tris-HCl, pH 8.5, 50 mM CaCl_2_, 50 mM BaCl_2_, and 32.5% PEG 400. All crystals were frozen in liquid nitrogen within 2 weeks and tested for X-ray diffraction at the Lawrence Berkeley National Laboratory ALS beamlines 5.0.1 (for the melibiose, α-MG, and α-NPG-containing complex datasets) or 5.0.2 (for the raffinose-containing datasets) using the remote data collection method.

ALS auto-processing XDS or DIALS programs output files were further reduction by AIMLESS in the ccp4i2 program for the structure solution (48). The statistics in data collection are described in **Table S1**.

### Structure determination

The structure determination was performed by the Molecular Replacement method using the α-NPG-bound D59C MelB_St_ mutant structure [PDB ID 7L17] as the search template, followed by rounds of manual building and refinement to resolutions of 2.60 Å, 3.05, 3.45, or 3.68 Å for structure with α-NPG, melibiose, raffinose, or α-MG, respectively, in Phenix (49). The model building and refinement were performed in Phenix and Coot (50), respectively. The structures were modeled from positions 2 to 453 or 455, respectively, without gaps, and the missing side chains due to density disorder, as well as the Ramachandran assessment, are listed in **Table S5**.

### Sugar docking and modeling

One strong positive density, with varying sizes and shapes, was observed in the difference maps of each of the four structures. The size and shape matched the sugars that co-crystallized, and the docked sugar molecules fitted well with the densities. The sugar refinement restraints were generated from SMILES using the ELBOW program in Phenix (49). To the 2.60 Å α-NPG-bound map, five water molecules were modeled. In addition, a PEG molecule (ligand ID 1PE) was also modeled to a strong positive density with a sausage shape aligning with helix IX. To the melibiose-bound structure, two water molecules were added.

### Hydrogen-deuterium exchange coupled to mass spectrometry (HDX-MS)

An in-solution HDX-MS experiment was performed to study the substrate-induced structural dynamics of MelB_St_. As described (21), the labeling, quenching, lipid removal, and online digestion were achieved using a fully automated manner using an HDx3 extended parallel system (LEAP Technologies, Morrisville, NC)(51, 52). MelB_St_ was prepared at 50.0 μM in a Na^+^-free buffer (25 mM Tris-HCl, pH 7.5, 150 mM choline chloride, 10% glycerol, 0.035% UDM in H_2_O), either in the absence of a substrate (apo) or in the presence of 100 mM melibiose, 100 mM NaCl, or both. The hydrogen/deuterium exchange reaction as described previously (21). Briefly, aliquots of 4 µl of each sample were diluted 10-fold into the labeling buffer (25 mM Tris-HCl, pD 7.5, 50 mM choline chloride, 10% glycerol, 0.035% UDM in D_2_O), without or with 100 mM of melibiose, Na^+^, or both. Labeled samples were incubated in D_2_O buffer at 20 °C for multiple time points (30 sec, 300 sec, and 3000 sec) in triplicate, and non-deuterated controls were prepared similarly, except that H_2_O buffer was used in the labeling step.

At each designated time point, reaction was quenched by adding an equal volume of ice-cold quench buffer (6 M urea, 100 mM citric acid, pH, 2.3 in H_2_O) for 180 seconds at 0 °C and immediately subjected to a lipid filtration module integrated on the LEAP PAL system. After incubation of a 60 sec with ZrO2 particles, the LEAP X-Press then compressed the filter assembly to separate proteins from the ZrO2 particles-bound phospholipids and detergents. The filtered protein sample was injected into a cool box for online digestion and separation.

LC/MS bottom-up HDX was performed using a Thermo Scientific™ Ultimate™ 3000 UHPLC system and Thermo Scientific^TM^ Orbitrap Eclipse^TM^ Tribrid^TM^ mass spectrometer. Samples were digested with a Nepenthesin-2 (Affipro, Czech Republic) column at 8 °C and then trapped in a 1.0 mm x 5.0 mm, 5.0 µm trap cartridge for desalting over 180 sec. The resulting peptides were then separated on a Thermo Scientific™ Hypersil Gold^TM^, 50 x1 mm, 1.9 µm, C18 column with a gradient of 10 % to 40 % gradient (A: water, 0.1 % formic acid; B: acetonitrile, 0.1 % formic acid) for 15 minutes at a flow rate of 40 µL/min. A pepsin wash was added in between runs to minimize the carryover.

A nonspecific digested peptide database has been created for MelB_St_ with a separate MS/MS measurement of non-deuterated samples as described (21). Digested peptides from undeuterated MelB_St_ protein were identified on the orbitrap mass spectrometer using the same LC gradient as the HDX-MS experiment. Using the Thermo BioPharma Finder software (v 5.1), MS2 spectra were matched to the MelB_St_ sequence with fixed modifications.

A total of 146 or 150 peptide assignments (with confident HDX data across all labeling times) were confirmed for MelB_St_ samples, resulting in 86-87% sequence coverage. The MS data were processed using the Sierra Analytics HDExaminer software with the MelB_St_ peptide database. Following the automated HDX-MS analysis, manual curation was performed. Upon the completion of the data review, a single charge state with high-quality spectra for all replicates across all HDX labeling times was chosen to represent HDX for each peptide. Differential HDX data were tested for statistical significance using the hybrid significance testing criteria method with an in-house MATLAB script, where the HDX differences at different protein states were calculated (ΔD = D_Holo_ - D_Apo_). Mean HDX differences from the three replicates were assigned as significant according to the hybrid criteria based on the pooled standard deviation and Welch’s t-test with P < 0.05. The statistically significant differences observed at each residue (ΔD_Mel-Apo_ < |0.186|, ΔD_(Na+)_ - Apo < |0.224|, or ΔD_Na(+)Mel_ - Apo < |0.175|) were used to map HDX consensus effects based on overlapping peptides onto the structure models.

### Statistics and reproducibility

All experiments were performed 2-4 times. The average values were presented in the table with standard errors. An unpaired t-test was used for statistical analysis. For the relative D%, the data were transferred to log values prior to the unpaired t-test.

### Graphs

Pymol (3.1.5.1) (53) and UCSF ChimeraX (X) were used to generate all graphs. The program Origin 2024 was used to plot the ITC curves and transport data.

### MD simulations

The crystal structure of the α-NPG-bound D59C MelB_St_ at a resolution of 2.6 Å was replaced with a melibiose molecule according to the melibiose-bound structure, and the D59C was mutated back to Asp. Both Asp59 and Asp55 residues were set in the deprotonated state. Three systems, including the apo, melibiose-bound, and melibiose- and Na^+^-bound states, were generated by embedding them in a POPE:POPG (7:2) lipid bilayer created using the CHARMM-GUI(54) web server. The lipid bilayer was capped with a 25 Å water box on each side, and ∼0.15 M NaCl was added to neutralize the system charge and mimic the ionic strength of physiological conditions. The resulting system has ∼120,000 atoms, with a periodic boundary condition of ∼110 × 110 × 125 (Å). The ff14SB(55), lipid17(56), and GLYCAM(57) force fields were employed to treat the protein, lipid, and melibiose, respectively. The TIP3P(58) water model was used for all water molecules.

After system assembly, energy minimization was carried out with harmonic restraints (1000 kJ/mol/Å²) on protein and lipid’s heavy atoms for 40,000 steps. This was followed by 200 ps of equilibration in the constant NVT ensemble at 300 K and 1 ns of NPT equilibration with gradually decreased force constants in the harmonic restraints. Five independent replicas of production simulations in the constant NPT ensemble were initiated from different snapshots in the NVT trajectory. For each replica, a trajectory was propagated for ∼400 ns at 300 K and 1 atm. The total sampling time for each system was ∼2 μs. The temperature was controlled with a Langevin thermostat(59) with a friction coefficient of 1 ps^-1^ and pressure with a Berendsen barostat(60) using anisotropic scaling with a relaxation time of 1 ps. Long-range electrostatics were treated with the Particle Mesh Ewald (PME) method(61) (tolerance 5 × 10^-4^), and van der Waals interactions were truncated using a 12 Å cutoff distance. A 2 fs timestep was used throughout all simulations. The SHAKE algorithm(62) was employed to constrain the lengths of all hydrogen-containing covalent bonds. All MD simulations were performed with the AMBER24 software package.(63)

Water-1 occupancy identified in the α-NPG-bound crystal structure was evaluated by monitoring interactions of water molecules with four coordinating residues. For each frame, a water molecule was defined as located in the sugar-binding pocket if the distances from its oxygen (O) atom to a few neighboring residues and the melibiose molecule satisfied the following rules. First, four pairs of heavy atoms on the melibiose molecule and neighboring residues were defined: O5 on the galactosyl ring and NE2 on Gln372, O5 and OG1 on Thr373, O6 and NE2 on Gln372, as well as O6 and OG1 on Thr373. Second, for each pair of the above-defined heavy atoms, the distance between the water oxygen atom (O) and each of the two heavy atoms in the pair was calculated. Third, out of all four pairs, if at least in one of them both distances were within 4 Å and at least one distance was below 3.5 Å, the water molecule was defined as occupying the sugar-binding site. The water-1 occupancy in the sugar-binding site was then computed as the fraction of frames containing at least one occupying water molecule over the total number of frames in all replicas of the trajectories.

The side-chain heavy-atom root-mean-square fluctuation (RMSF) of all residues in the sugar- and cation-binding pockets were determined using CPPTRAJ.

## Data availability statement

The x-ray diffraction datasets and models have been deposited to wwPDB under the accession codes 9OLD for the α-nitrophenyl galactoside-bound, 9OLI for the melibiose-bound, 9OLR for the α-methyl galactoside, as well as 9OLP for the raffinose-bound D59C MelB_St_.

## Acknowledgments

The authors thank Dr. William Mallard for creating Fig. 5a. The X-ray diffraction datasets were collected at ALS BL 5.01 or 5.0.2. This work was supported by the National Institutes of Health Grants R35GM153222 to L.G. and R35GM150780 to R.L.

## Abbreviations

MelB_St_: *Salmonella enterica* serovar Typhimurium melibiose permease
MFS: major facilitator superfamilies
ITC: isothermal titration calorimetry
α-NPG: *p*-nitrophenyl-α-D-galactopyranoside
Mel: melibiose
α-MG: methyl α-D-galactoside
Raff: raffinose
*K*_d_: dissociation constant
HDX-MS: hydrogen-deuterium exchange coupled to mass spectrometry
D: deuterium
RMSF: root-mean-square fluctuation

## References

1. Ferrada, E., and Superti-Furga, G. (2022) A structure and evolutionary-based classification of solute carriers. iScience 25, 105096

2. Pao, S. S., Paulsen, I. T., and Saier, M. H., Jr. (1998) Major facilitator superfamily. Microbiol Mol Biol Rev 62, 1–32

3. Cesar-Razquin, A., Snijder, B., Frappier-Brinton, T., Isserlin, R., Gyimesi, G., Bai, X. et al. (2015) A Call for Systematic Research on Solute Carriers. Cell 162, 478–487

4. Lin, L., Yee, S. W., Kim, R. B., and Giacomini, K. M. (2015) SLC transporters as therapeutic targets: emerging opportunities. Nat Rev Drug Discov 14, 543–560

5. Guan, L., and Kaback, H. R. (2006) Lessons from lactose permease. Annu Rev Biophys Biomol Struct 35, 67–91

6. Yan, N. (2015) Structural Biology of the Major Facilitator Superfamily Transporters. Annu Rev Biophys 44, 257–283

7. Drew, D., North, R. A., Nagarathinam, K., and Tanabe, M. (2021) Structures and General Transport Mechanisms by the Major Facilitator Superfamily (MFS). Chem Rev 121, 5289–5335

8. Guan, L. (2023) The rapid developments of membrane protein structure biology over the last two decades. BMC Biol 21, 300

9. Nguyen, L. N., Ma, D., Shui, G., Wong, P., Cazenave-Gassiot, A., Zhang, X. et al. (2014) Mfsd2a is a transporter for the essential omega-3 fatty acid docosahexaenoic acid. Nature 509, 503–506

10. Cater, R. J., Chua, G. L., Erramilli, S. K., Keener, J. E., Choy, B. C., Tokarz, P. et al. (2021) Structural basis of omega-3 fatty acid transport across the blood-brain barrier. Nature 595, 315–319

11. Chua, G. L., Tan, B. C., Loke, R. Y. J., He, M., Chin, C. F., Wong, B. H. et al. (2023) Mfsd2a utilizes a flippase mechanism to mediate omega-3 fatty acid lysolipid transport. Proc Natl Acad Sci U S A 120, e2215290120

12. Wilson, T. H., and Ding, P. Z. (2001) Sodium-substrate cotransport in bacteria. Biochim Biophys Acta 1505, 121–130

13. Maehrel, C., Cordat, E., Mus-Veteau, I., and Leblanc, G. (1998) Structural studies of the melibiose permease of *Escherichia coli* by fluorescence resonance energy transfer. I. Evidence for ion-induced conformational change. J Biol Chem 273, 33192–33197

14. Meyer-Lipp, K., Sery, N., Ganea, C., Basquin, C., Fendler, K., and Leblanc, G. (2006) The inner interhelix loop 4-5 of the melibiose permease from *Escherichia coli* takes part in conformational changes after sugar binding. J Biol Chem 281, 25882–25892

15. Guan, L., Nurva, S., and Ankeshwarapu, S. P. (2011) Mechanism of melibiose/cation symport of the melibiose permease of *Salmonella typhimurium*. J Biol Chem 286, 6367–6374

16. Granell, M., Leon, X., Leblanc, G., Padros, E., and Lorenz-Fonfria, V. A. (2010) Structural insights into the activation mechanism of melibiose permease by sodium binding. Proc Natl Acad Sci U S A 107, 22078–22083

17. Guan, L. (2018) Na(+)/Melibiose Membrane Transport Protein, MelB In In: Roberts G., Watts A., European Biophysical Societies (eds) Encyclopedia of Biophysics, 2nd Ed. Roberts G, Watts A, eds. Springer, Berlin, Heidelberg, 10.1007/978-3-642-35943-9_10082-1

18. Ethayathulla, A. S., Yousef, M. S., Amin, A., Leblanc, G., Kaback, H. R., and Guan, L. (2014) Structure-based mechanism for Na(+)/melibiose symport by MelB. Nat Commun 5, 3009

19. Hariharan, P., and Guan, L. (2017) Thermodynamic cooperativity of cosubstrate binding and cation selectivity of Salmonella typhimurium MelB. J Gen Physiol 149, 1029–1039

20. Guan, L., and Hariharan, P. (2021) X-ray crystallography reveals molecular recognition mechanism for sugar binding in a melibiose transporter MelB. Commun Biol 4, 931

21. Hariharan, P., Shi, Y., Katsube, S., Willibal, K., Burrows, N. D., Mitchell, P. et al. (2024) Mobile barrier mechanisms for Na(+)-coupled symport in an MFS sugar transporter. Elife 12,

22. Hariharan, P., Bakhtiiari, A., Liang, R., and Guan, L. (2024) Distinct roles of the major binding residues in the cation-binding pocket of the melibiose transporter MelB. J Biol Chem 107427

23. Katsube, S., Liang, R., Amin, A., Hariharan, P., and Guan, L. (2022) Molecular Basis for the Cation Selectivity of Salmonella typhimurium Melibiose Permease. J Mol Biol 434, 167598

24. Guan, L., Jakkula, S. V., Hodkoff, A. A., and Su, Y. (2012) Role of Gly117 in the cation/melibiose symport of MelB of *Salmonella typhimurium*. Biochemistry 51, 2950–2957

25. Hariharan, P., and Guan, L. (2014) Insights into the inhibitory mechanisms of the regulatory protein IIA(Glc) on melibiose permease activity. J Biol Chem 289, 33012–33019

26. Liang, R., and Guan, L. (2024) Atomic-Level Free Energy Landscape Reveals Cooperative Symport Mechanism of Melibiose Transporter. eLife https://doiorg/107554/eLife103421

27. Hamuro, Y. (2024) Interpretation of Hydrogen/Deuterium Exchange Mass Spectrometry. J Am Soc Mass Spectrom 35, 819–828

28. Masson, G. R., Burke, J. E., Ahn, N. G., Anand, G. S., Borchers, C., Brier, S. et al. (2019) Recommendations for performing, interpreting and reporting hydrogen deuterium exchange mass spectrometry (HDX-MS) experiments. Nat Methods 16, 595–602

29. Wilson, D. M., and Wilson, T. H. (1987) Cation specificity for sugar substrates of the melibiose carrier in Escherichia coli. Biochim Biophys Acta 904, 191–200

30. Hariharan, P., and Guan, L. (2021) Cooperative binding ensures the obligatory melibiose/Na+ cotransport in MelB. J Gen Physiol 153,

31. Amin, A., Ethayathulla, A. S., and Guan, L. (2014) Suppression of conformation-compromised mutants of *Salmonella enterica* serovar Typhimurium MelB. J Bacteriol 196, 3134–3139

32. Jia, R., Martens, C., Shekhar, M., Pant, S., Pellowe, G. A., Lau, A. M. et al. (2020) Hydrogen-deuterium exchange mass spectrometry captures distinct dynamics upon substrate and inhibitor binding to a transporter. Nat Commun 11, 6162

33. Zmyslowski, A. M., Baxa, M. C., Gagnon, I. A., and Sosnick, T. R. (2022) HDX-MS performed on BtuB in E. coli outer membranes delineates the luminal domain’s allostery and unfolding upon B12 and TonB binding. Proc Natl Acad Sci U S A 119, e2119436119

34. Hageman, T. S., and Weis, D. D. (2019) Reliable Identification of Significant Differences in Differential Hydrogen Exchange-Mass Spectrometry Measurements Using a Hybrid Significance Testing Approach. Anal Chem 91, 8008–8016

35. Markham, K. J., Tikhonova, E. B., Scarpa, A. C., Hariharan, P., Katsube, S., and Guan, L. (2021) Complete cysteine-scanning mutagenesis of the Salmonella typhimurium melibiose permease. J Biol Chem 297, 101090

36. Jakkula, S. V., and Guan, L. (2012) Reduced Na(+) affinity increases turnover of *Salmonella enterica* serovar Typhimurium MelB. J Bacteriol 194, 5538–5544

37. Blaimschein, N., Hariharan, P., Manioglu, S., Guan, L., and Muller, D. J. (2023) Substrate-binding guides individual melibiose permeases MelB to structurally soften and to destabilize cytoplasmic middle-loop C3. Structure 31, 58–67 e54

38. Smirnova, I. N., Kasho, V., and Kaback, H. R. (2008) Protonation and sugar binding to LacY. Proc Natl Acad Sci U S A 105, 8896–8901

39. Grytsyk, N., Sugihara, J., Kaback, H. R., and Hellwig, P. (2017) pKa of Glu325 in LacY. Proc Natl Acad Sci U S A 114, 1530–1535

40. Guan, L., Mirza, O., Verner, G., Iwata, S., and Kaback, H. (2007) Structural determination of wild-type lactose permease. Proceedings of the National Academy of Sciences of the United States of America 104, 15294–15298

41. Mirza, O., Guan, L., Verner, G., Iwata, S., and Kaback, H. R. (2006) Structural evidence for induced fit and a mechanism for sugar/H+ symport in LacY. EMBO J 25, 1177–1183

42. Kumar, H., Kasho, V., Smirnova, I., Finer-Moore, J. S., Kaback, H. R., and Stroud, R. M. (2014) Structure of sugar-bound LacY. Proc Natl Acad Sci U S A 111, 1784–1788

43. Sun, L., Zeng, X., Yan, C., Sun, X., Gong, X., Rao, Y. et al. (2012) Crystal structure of a bacterial homologue of glucose transporters GLUT1-4. Nature 490, 361–366

44. Jia, R., Bradshaw, R. T., Calvaresi, V., and Politis, A. (2023) Integrating Hydrogen Deuterium Exchange-Mass Spectrometry with Molecular Simulations Enables Quantification of the Conformational Populations of the Sugar Transporter XylE. J Am Chem Soc 145, 7768–7779

45. Smirnova, I., Kasho, V., and Kaback, H. R. (2011) Lactose permease and the alternating access mechanism. Biochemistry 50, 9684–9693

46. Botfield, M. C., and Wilson, T. H. (1988) Mutations that simultaneously alter both sugar and cation specificity in the melibiose carrier of Escherichia coli. J Biol Chem 263, 12909–12915

47. Pourcher, T., Leclercq, S., Brandolin, G., and Leblanc, G. (1995) Melibiose permease of Escherichia coli: large scale purification and evidence that H(+), Na(+), and Li(+) sugar symport is catalyzed by a single polypeptide. Biochemistry 34, 4412–4420

48. Potterton, L., Agirre, J., Ballard, C., Cowtan, K., Dodson, E., Evans, P. R. et al. (2018) CCP4i2: the new graphical user interface to the CCP4 program suite. Acta Crystallogr D Struct Biol 74, 68–84

49. Liebschner, D., Afonine, P. V., Baker, M. L., Bunkoczi, G., Chen, V. B., Croll, T. I. et al. (2019) Macromolecular structure determination using X-rays, neutrons and electrons: recent developments in Phenix. Acta Crystallogr D Struct Biol 75, 861–877

50. Casanal, A., Lohkamp, B., and Emsley, P. (2020) Current developments in Coot for macromolecular model building of Electron Cryo-microscopy and Crystallographic Data. Protein Sci 29, 1069–1078

51. Hamuro, Y., Coales, S. J., Southern, M. R., Nemeth-Cawley, J. F., Stranz, D. D., and Griffin, P. R. (2003) Rapid analysis of protein structure and dynamics by hydrogen/deuterium exchange mass spectrometry. J Biomol Tech 14, 171–182

52. Hamuro, Y., and Coales, S. J. (2018) Optimization of Feasibility Stage for Hydrogen/Deuterium Exchange Mass Spectrometry. J Am Soc Mass Spectrom 29, 623–629

53. Schrodinger, L. (2013) The PyMOL Molecular Graphics System, Version 1.5.

54. Wu, E. L., Cheng, X., Jo, S., Rui, H., Song, K. C., Dávila Contreras, E. M. et al. (2014) CHARMM GUI membrane builder toward realistic biological membrane simulations Wiley Online Library,

55. Maier, J. A., Martinez, C., Kasavajhala, K., Wickstrom, L., Hauser, K. E., and Simmerling, C. (2015) ff14SB: improving the accuracy of protein side chain and backbone parameters from ff99SB. Journal of chemical theory and computation 11, 3696–3713

56. Skjevik, Å. A., Madej, B. D., Dickson, C. J., Lin, C., Teigen, K., Walker, R. C. et al. (2016) Simulation of lipid bilayer self-assembly using all-atom lipid force fields. Phys Chem Chem Phys 18, 10573–10584

57. Group, W. (2005-2023) GLYCAM Web Complex Carbohydrate Research Center, University of Georgia, Athens, GA (http://glycam.org)

58. Jorgensen, W. L., Chandrasekhar, J., Madura, J. D., Impey, R. W., and Klein, M. L. (1983) Comparison of simple potential functions for simulating liquid water. J Chem Phys 79, 926–935

59. Sindhikara, D. J., Kim, S., Voter, A. F., and Roitberg, A. E. (2009) Bad seeds sprout perilous dynamics: Stochastic thermostat induced trajectory synchronization in biomolecules. Journal of Chemical Theory and Computation 5, 1624–1631

60. Uberuaga, B. P., Anghel, M., and Voter, A. F. (2004) Synchronization of trajectories in canonical molecular-dynamics simulations: Observation, explanation, and exploitation. J Chem Phys 120, 6363–6374

61. Darden, T., York, D., and Pedersen, L. (1993) Particle mesh Ewald: An N log (N) method for Ewald sums in large systems. J Chem Phys 98, 10089–10089

62. Ryckaert, J.-P., Ciccotti, G., and Berendsen, H. J. (1977) Numerical integration of the cartesian equations of motion of a system with constraints: molecular dynamics of n-alkanes. J Comput Phys 23, 327–341

63. Case, D. A., Aktulga, H. M., Belfon, K., Ben-Shalom, I. Y., Berryman, J. T., Brozell, S. R. et al. (2023) Amber 2023, University of California, San Francisco,

