## Supplemental Table S1 for "Allosteric effects of the coupling cation in melibiose transporter MelB"

**Table S1. Crystallographic Data collection, phase, and refinement statistics**

| **Data collection** | D59C $MelB_{St}$ complexed with α-galactosides | | | |
|---|---|---|---|---|
| | α-NPG [PDB ID, 9OLD] | Melibiose [PDB ID, 9OLI] | α-Methyl galactoside [PDB ID, 9OLR] | Raffinose [PDB ID, 9OLP] |
| | ALS 5.0.2 | ALS 5.0.2 | ALS 5.0.2 | ALS 5.0.1 |
| | 0.9795 | 0.9795 | 0.9795 | 0.97 |
| Space group | P 31 2 1 | P 31 2 1 | P 31 2 1 | P 31 2 1 |
| Cell dimensions | | | | |
| *a*, *b*, *c* (Å) | 126.9 126.9 104.4 | 126.4 126.4 103.5 | 127.496 127.496 106.039 | 127.1 127.1 104.9 |
| α, β, γ (°) | 90 90 120 | 90 90 120 | 90 90 120 | 90 90 120 |
| Resolution (Å) | 20 - 2.60 | 20 - 3.05 | 20 - 3.68 | 20 - 3.40 |
| $R_{meas}$ | 0.072 (3.299) | 0.067 (2.064) | 0.145 (0.508) | 0.339 (1.297) |
| $I / \sigma I$ | 16.5 (0.9) | 11.8 (0.8) | 8.4 (4.4) | 7.9 / 3.1 |
| CC(1/2) | | 1.00 (0.26) | 1.00 (0.93) | 1.00 (0.873) |
| Completeness (%) | 98.6 (98.4) | 99.50 (100) | 99.8 (93.3) | 99.5 (100) |
| Redundancy | 9.7 (10.3) | 5.7 (5.7) | 9.2 (8.8) | 18.3 (17.5) |
| **Refinement** | | | | |
| Resolution (Å) | 20 - 2.60 (2.63 - 2.60) | 20 - 3.05 (3.13 - 3.05) | 20 - 3.68 (4.05 - 3.68) | 20 - 3.40 (3.54 - 3.40) |
| No. reflections | 29768 (1617) | 18439 (1410) | 10983 (2671) | 13237 (1424) |
| $R_{work}$ / $R_{free}$ | 0.256/0.287 | 0.255 / 0.294 | 0.297 / 0.330 | 0.316 / 0.329 |
| No. atoms | 3584 | 3540 | 3570 | 3581 |
| Protein | 3542 | 3515 | 3557 | 3547 |
| Ligand/ion | 37 | 23 | 13 | 34 |
| Water | 5 | 2 | 0 | 0 |
| *B*-factors | 121.88 | 135.70 | 116.76 | 113.7 |
| Protein | 121.93 | 135.64 | 116.78 | 113.4 |
| Ligand/ion | 121.31 | 146.75 | 109.95 | 109.6 |
| Water | 95.54 | 109.73 | / | / |
| R.m.s. deviations | | | | |
| Bond lengths (Å) | 0.003 | 0.003 | 0.002 | 0.002 |
| Bond angles (°) | 0.573 | 0.545 | 0.486 | 0.565 |

A single crystal was used for all structures. Values in parentheses are for highest-resolution shell.
