## Supplemental Table S2 for "Allosteric effects of the coupling cation in melibiose transporter MelB"

**Table S2. HDX reaction, labeling details, and statistics**

| | $\Delta D_{Mel - Apo}$ | $\Delta D_{Na(+) - Apo}$ | $\Delta D_{Na(+)Mel – Apo}$# |
|---|---|---|---|
| Samples measured | Test-1: WT $MelB_{St}$ (Apo);<br>Test-2: WT $MelB_{St}$ with 50 mM melibiose. | Test-3: WT $MelB_{St}$ (Apo);<br>Test-4: WT $MelB_{St}$ with 150 mM $Na^+$. | Test-5: WT $MelB_{St}$ with 50 mM melibiose and 150 mM $Na^+$. |
| HX reaction buffer | 25 mM Tris-HCl, pD 7.5, 150 mM NaCl, 10% Glycerol, and 0.01% DDM | 25 mM Tris-HCl, pD 7.5, 150 mM NaCl, 10% Glycerol, and 0.01% DDM | 25 mM Tris-HCl, pD 7.5, 150 mM NaCl, 10% Glycerol, and 0.01% DDM |
| Reaction temperature (°C) | 20 | 20 | 20 |
| HX time course (s) | 0, 30, 300, 3000 | 0, 30, 300, 3000 | 0, 30, 300, 3000 |
| Number of peptides | 150 | 152 | 150 |
| Sequence coverage by labeling | 87.47 | 86.62 | 87.47 |
| Mean peptide length | 8.4 | 7.9 | 8.0 |
| Average redundancy | 3.7 | 3.5 | 3.5 |
| Replicates (technical) | 3 | 3 | 3 |
| $\|\Delta D\|$ (Da) | 0.186 | 0.224 | 0.175 |
| Back exchange rate | Not appliable | Not appliable | Not appliable |
| Number of non-covered positions | 59 | 63 | 59 |
| Threshold | ± 0.186 | ± 0.224 | ± 0.175 |
| Number of overlapping peptides with significant ΔD > \| Threshold \| and P < 0.05 | 27 | 21 | 30 |
| Number of covered residues with significant ΔD > \| Threshold \| and P < 0.05 | 153 | 122 | 133 |
| Number of covered residues with insignificant ΔD <\| Threshold \| and P > 0.05 | 237 | 264 | 257 |

### Test-1 data of Apo $MelB_{St}$ was used for comparison and calculation.
