## Supplemental Table S3 for "Allosteric effects of the coupling cation in melibiose transporter MelB"

### Table S3. Relative deuterium uptake and uncovered positions of the Apo $MelB_{St}$

| Region | Fragment (#residues) | HDX coverage (# residues) | Averaged D%* (n = 2) | 59 of uncovered fragments (#residues) | Sequence |
|---|---|---|---|---|---|
| H-I | 4 - 38 (35) | 2 - 35 (34) | 8.531 ± 0.865# | 0 | |
| H-II | 40 - 68 (29) | 36 - 77 (42) | 1.736 ± 0.345 | 0 | |
| H-III | 76 - 100 (25) | 76 - 100 (25) | 1.465 ± 0.037 | Position 92 (1) | $F^{92}$ |
| H-IV | 103 - 135 (33) | 99 - 138 (40) | 2.083 ± 0.004 | Positions 111-119 (9) | $V^{111}TYILWGMT^{119}$ |
| H-V | 137 - 171 (35) | 137 - 169 (32) | 7.069 ± 0.022# | 0 | |
| H-VI | 174 - 201 (28) | 170 - 205 (36) | 1.627 ± 0.205 | Positions 181-183 (3) | $F^{182}TL$ |
| $Loop_{6-7}$ | 202 - 230 (29) | 209 - 233 (25) | 4.955 ± 0.291 | Positions 206-208 (3) | $S^{206}SD$ |
| H-VII | 231 - 261 (31) | 227 - 263 (37) | 1.767 ± 0.136 | 0 | |
| H-VIII | 266 - 285 (20) | 264 - 283 (20) | 3.587 ± 0.218 | Positions 272-273 (2) | $L^{272}S^{273}$ |
| $Loop_{8-9}$ | 286 - 295 (10) | 282 - 298 (17) | 10.876 ± 0.189 | 0 | |
| H-IX | 296 - 319 (24) | 296 - 314 (18) | 1.921 ± 0.037 | Positions 303-308, 315-316 (8) | $S^{303}VMPVL^{308}$ $A^{315}M^{316}$ |
| $Loop_{9-10}$ | 320-323 (4) | 317-326 (10) | 6.475 ± 1.165 | 0 | |
| H-X^ | 323 - 360 (38) | 342 - 360 (18)## | 1.992 ± 0.388 | Positions 327-341 and 345-348 (19) | $I^{327}VAAGIFLNIGT ALF^{341}$ $Q^{345}VIM^{348}$ |
| $Loop_{10-11}$ | 361 – 365 (6) | 364 - 368 (5) | 19.94 | Positions 360-363 (4) | $L^{360}NIR^{363}$ |
| H-XI | 366 – 395 (30) | 369 - 386 (19) | 2.207 ± 0.007 | Positions 387-393 (7) | $I^{387}ALVLGL^{393}$ |
| $Loop_{11-12}$^ | 396 – 403 (8) | 394 - 406 (13)## | 3.581 ± 2.046 | 0 | |
| H-XII | 404 – 432 (31) | 406 - 435 (30) | 1.395 ± 0.184 | 0 | |
| C-term Tail^ | 433 – 476 (38) | 439 - 470 (32)## | 40.890 ± 3.507 | Positions 436 – 438 (3) | $N^{436}GD^{438}$ |

*, Average from two dataset of apo state of the mean values of relative deuterium uptake (D%) across all time points (30 sec, 300 sec, and 3000 sec) of all covered peptides.

^, The uncovered residues from the dataset of $\Delta D_{Na(+)-Apo}$ are different, as listed here: H-X (Positions 327-348; 22 residues); $Loop_{10-11}$ (Positions 360-363; 3 residues); $Loop_{12-CTH}$ (Positions 436-440; 5 residues). The total number of the uncovered positions for Na+ vs. apo dataset is 63.

#, Averaged D% of helix I vs helix II, P =0.01; D% of helix V vs helix II, P =0.001. Unpaired t-test was applied for the data after log transformation.

##, the starting or sending positions between the two datasets were slightly different, with positions 349-360, 394-407 and 441-470 presented in the apo date from the $\Delta D_{Na(+)-Apo}$ dataset, respectively.
