## Supplemental Table S4 for "Allosteric effects of the coupling cation in melibiose transporter MelB"

**Table S4. HDX results at the sugar- and $Na^+$-binding pockets**

| | Mel (0.1857*) | | | $Na^+$ (0.224) | | | Mel with $Na^+$ (0.1754) | | |
|---|---|---|---|---|---|---|---|---|---|
| | Peptides | Data (P<0.05) | Protection^ | Peptid es | Data (P<0.05)$ | Protection^ | Peptid es | Data (P<0.05)$ | Protection^ |
| **Sugar-binding residues** | | | | | | | | | |
| K18 | 4 | 2 | 1 | 3 | 2 | 1 | 3 | 3 | 2 |
| D19 | 3 | 2 | 1 | 4 | 3 | 2 | 3 | 3 | 2 |
| I22 | 1 | 1 | 1 | 4 | 4 | 2 | 1 | 1 | 1 |
| Y26 | 3 | 1 | 1 | 3 | 3 | 1 | 3 | 5 | 1 |
| Y120 | 1 | 1 | 0 | 1 | 0 | / | 1 | 2 | 0 |
| D124 | 2 | 1 | 0 | 2 | 3 | 0 | 2 | 2 | 0 |
| W128 | 2 | 0 | / | 3 | 5 | 0 | 3 | 3 | 0 |
| R149 | 9 | 17 | 8 | 9 | 17 | 7 | 9 | 27 | 18 |
| A152 | 1 | 1 | 0 | 1 | 0 | 0 | 1 | 2 | 1 |
| W342 | 1 | 0 | / | 0 | / | / | 1 | 0 | / |
| Q372 | 1 | 2 | 0 | 1 | 3 | 1 | 1 | 3 | 1 |
| T373 | 2 | 2 | 0 | 2 | 3 | 1 | 2 | 3 | 1 |
| V376 | 2 | 0 | / | 3 | 2 | 0 | 2 | 1 | 0 |
| K377 | 2 | 0 | / | 3 | 2 | 0 | 2 | 1 | 0 |
| **Cation-binding residues** | | | | | | | | | |
| 55 | 3 | 2 | 0 | 3 | 5 | 0 | 3 | 2 | 0 |
| 58 | 3 | 2 | 0 | 3 | 5 | 0 | 3 | 2 | 0 |
| 59 | 4 | 2 | 0 | 4 | 5 | 0 | 4 | 2 | 0 |
| 121 | 2 | 1 | 0 | 2 | 1 | 0 | 2 | 3 | 0 |

* Threshold values.

^ $P < 0.05$ and D > | threshold | at any time point.
