## Supplemental Table S5 for "Allosteric effects of the coupling cation in melibiose transporter MelB"

**Table S5. Structure information**

| PDB ID (Ligand) | 9OLD (α-NPG) | 9OLI (Melibiose) | 9OLR (α-MG) | 9OLP (Raffinose) |
|---|---|---|---|---|
| Resolved positions | 2-255 | 2-253 | 2-254 | 2-254 |
| Missing side chains | Lys221<br>Lys291<br>His322<br>Arg431<br>Lys450<br>Lys453 | Arg70<br>Arg199<br>Val261<br>Leu267<br>Lys291<br>Asp320<br>His322<br>Leu334<br>Asn399<br>Lys450 | Lys291 | Arg70<br>His322<br>Leu447 |
| Ramachandran<br>Favored (%)<br>Outliers<br><br>Clash scores | <br>97.57<br>0.00<br><br>1.25 | <br>96.89<br>0.22<br><br>1.40 | <br>94.46<br>1.33<br><br>2.9 | <br>94.46<br>0.89<br><br>4.42 |
