## Supplemental Table S6 for "Allosteric effects of the coupling cation in melibiose transporter MelB"

**Table S6.** MD simulations of Wat-1 occupancy in sugar-bound $MelB_{St}$ with or without $Na^+$

| System | Replica | Occupancy |
|---|---|---|
| **Sugar + $Na^+$** | 1 | 97.49% |
| | 2 | 92.16% |
| | 3 | 98.86% |
| | 4 | 96.48% |
| | 5 | 99.21% |
| | **Average** | **96.84% (± 2.83%^)** |
| **Sugar Only** | 1 | 99.25% |
| | 2 | 94.64% |
| | 3 | 99.56% |
| | 4 | 96.42% |
| | 5 | 94.31% |
| | **Average** | **96.84% (± 2.48%)** |

**^, SD**
