## Supplemental Figs. 1-6 for "Allosteric effects of the coupling cation in melibiose transporter MelB"

**Fig S1**

**Superposition of four sugar-bound crystal structures of D59C $MelB_{St}$**

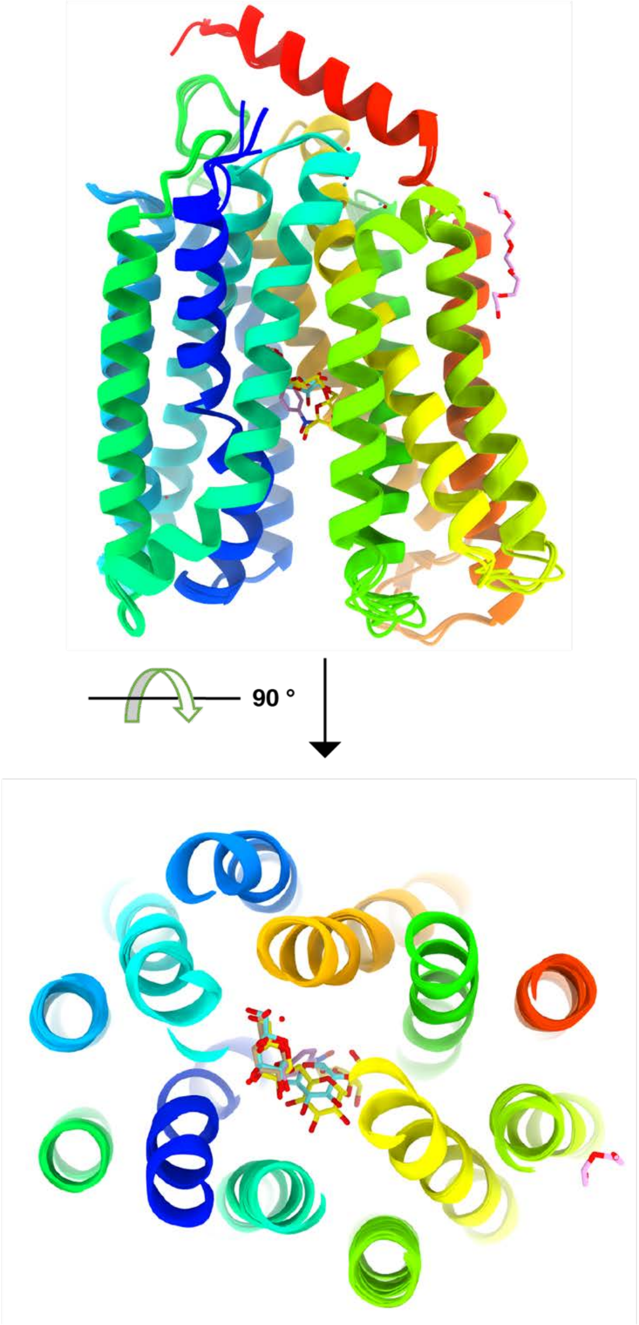


**Fig. S1. Overlay.** Four crystal structures of the D59C $MelB_{St}$ with melibiose, $\alpha$-MG, raffinose and $\alpha$-NPG bound, respectively, were superimposed with the MRSD values < 0.4 Å. Top, side view with a rainbow color code from N- to C-termini. Bottom, viewed from the cytoplasmic side.

**Fig. S2**

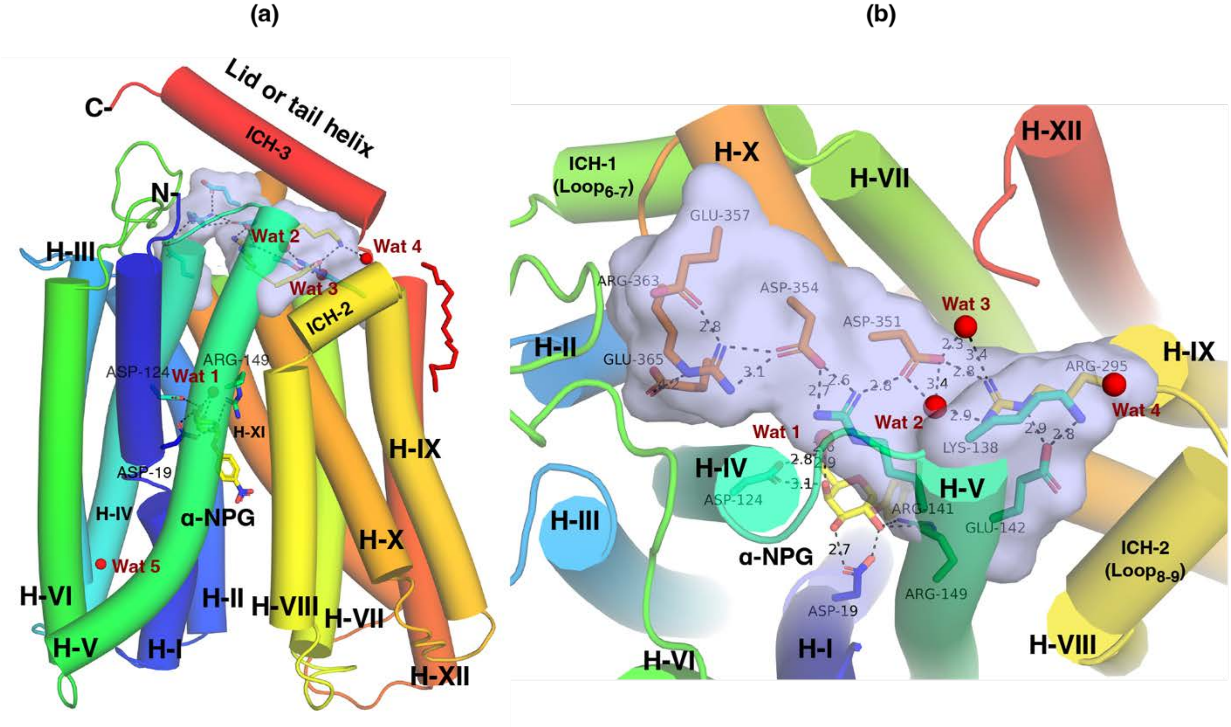


**Fig. S2. Waters interacting with the salt-bridge network.** Nine charged residues, including the three N-terminal residues (Lys138, Arg141, and Glu142 on helix V) colored in yellow and the six C-terminal residues (Arg295, Asp351, Asp354, Glu357, Rrg363, and Glu365 on helix XI and Loop$_{10\text{-}11}$) colored in cyan, are presented in sticks and highlighted in surface representation in light blue. Red sphere, water molecule. Wat 1 is bound with α-NPG and Thr373; Wat 2 and Wat 3 is associate with the salt-bridge pair Arg295 and Asp351, respectively; and Wat 4 is interacting with Lye138. The α-NPG is colored yellow and labeled. The transmembrane helices are labeled in roman numerical. (**a**) Side view; **(b)** Top view.

Fig. S3

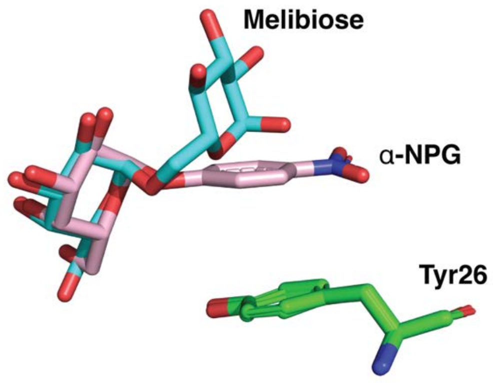


**Fig. S3**. **Overlay of the bound melibiose and α-NPG.** The crystal structures of D59C $MelB_{St}$ with melibiose or α-NPG bound were aligned, and the bound substrates were highlighted. The Tyr26 on helix I stacks with the phenyl ring of α-NPG.

Fig. S4

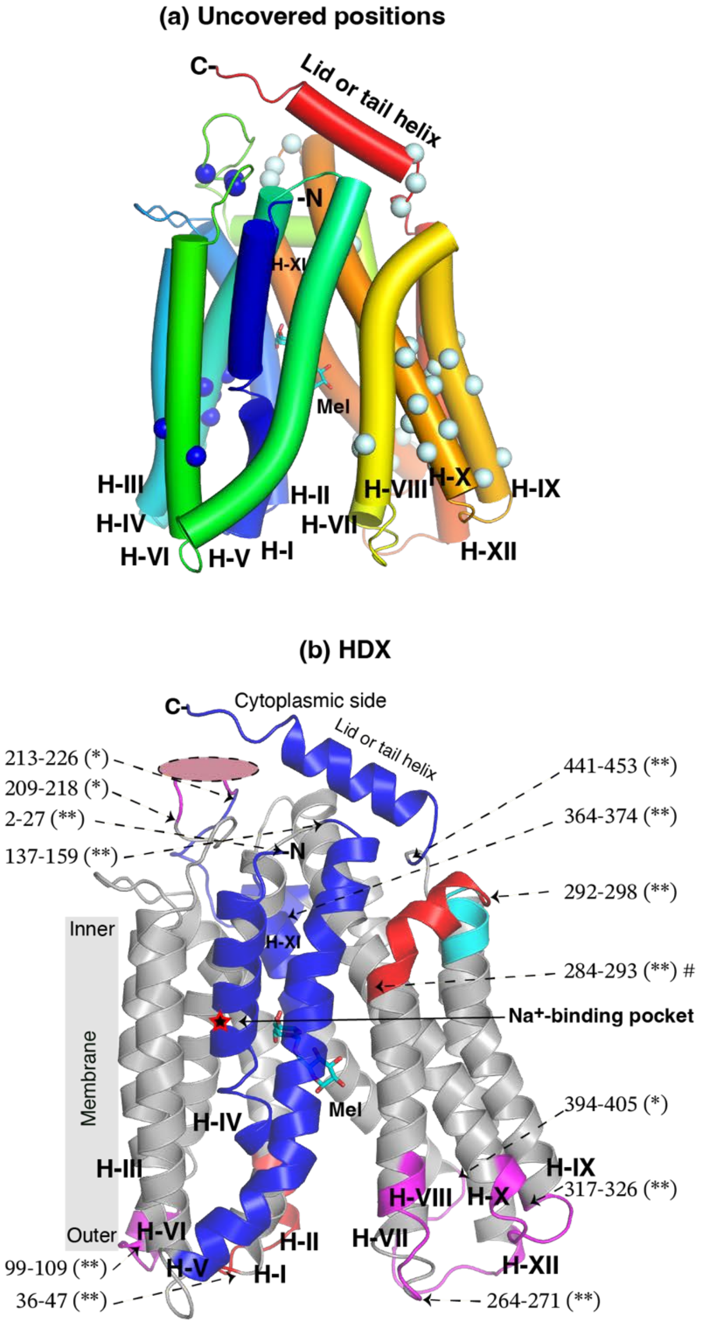


**Blue**, protected by Mel, $Na^+$, or Mel+$Na^+$
**Red**, deprotected by Mel or Mel+$Na^+$
**Magenta**, deprotected by Mel; insignificant ΔD or P>0.05 for $Na^+$ and Mel+$Na^+$
**Cyan**, deprotected by Mel but protected by Mel+$Na^+$
#, deprotected by Mel+$Na^+$ (P<0.001), and deprotectd by Mel (P=0.15)
(*), ΔD values are statistically significant, P<0.05
(**), ΔD values are statistically highly significant, P<0.01

**Fig. S4**. **Mapping of HDX data on an inward-facing melibiose-bound D59C $\mathrm{MelB_{St}}$. (a) Non-covered residues.** Transmembrane helices are labeled in Roman numerals. The non-covered positions on the N-terminal domains and C-terminal domain including extended loops are showed in α-carbon position and colored in blue and pale cyan, respectively. **(b)** Overlapping peptide with ligand-induced protection and deprotection of deuterium uptakes. Peptides with ΔD value greater than the absolute value of the threshold and $P < 0.05$ at any time point were colored according to the legend on the figure. The membrane region of $\mathrm{MelB_{St}}$ is indicated by a gray bar. Each peptide or overlapping peptide was indicated by arrows pointing to the starting position of the peptide.

**Fig. S5a. Deuterium uptake in the absence or presence of melibiose**

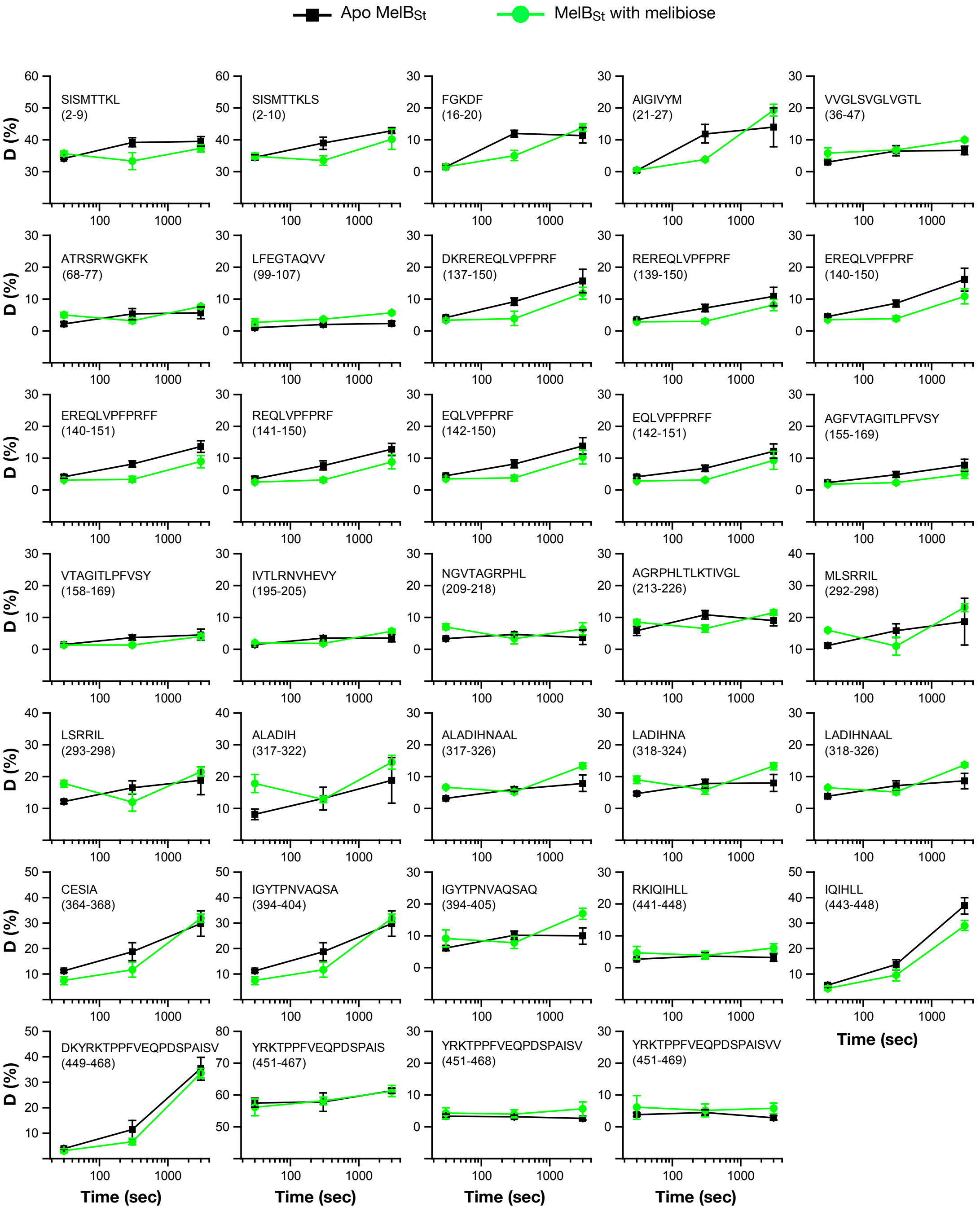

**Fig. S5b-1. Deuterium uptake in the absence or presence of Na+**

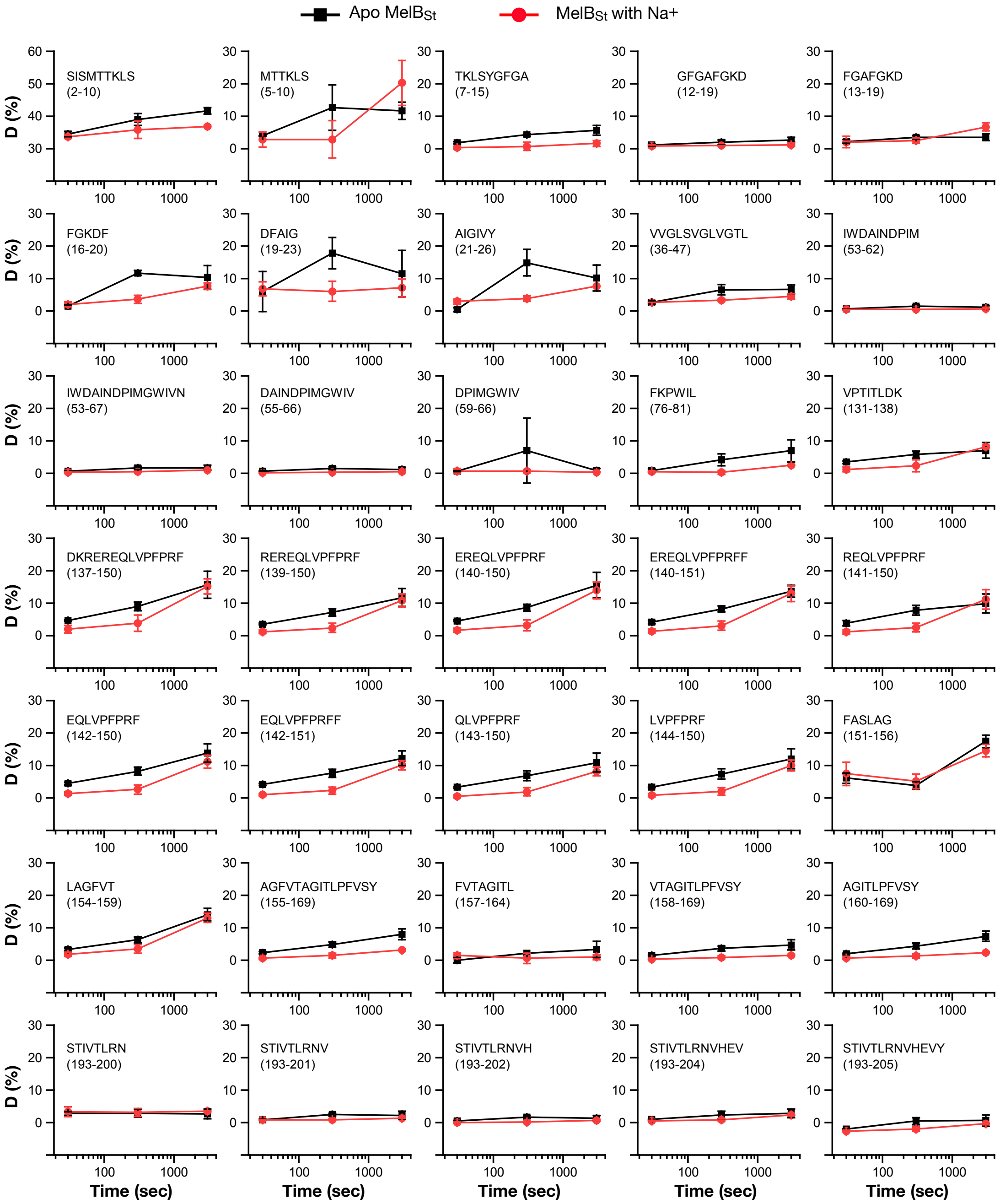

**Fig. S5b-2. Deuterium uptake in the absence or presence of Na+**

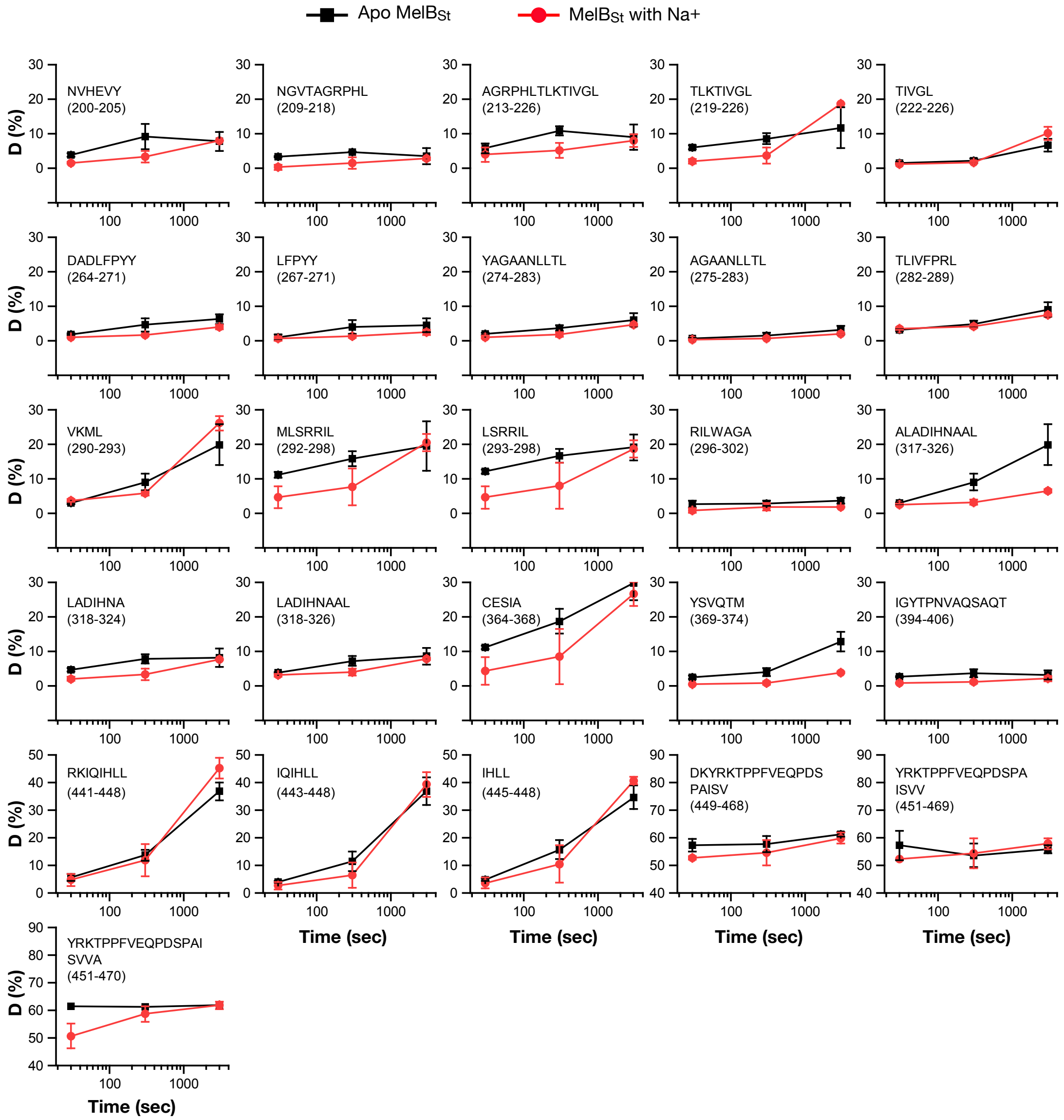

**Fig. S5c-1. Deuterium uptake in the absence or presence of melibiose and Na+**

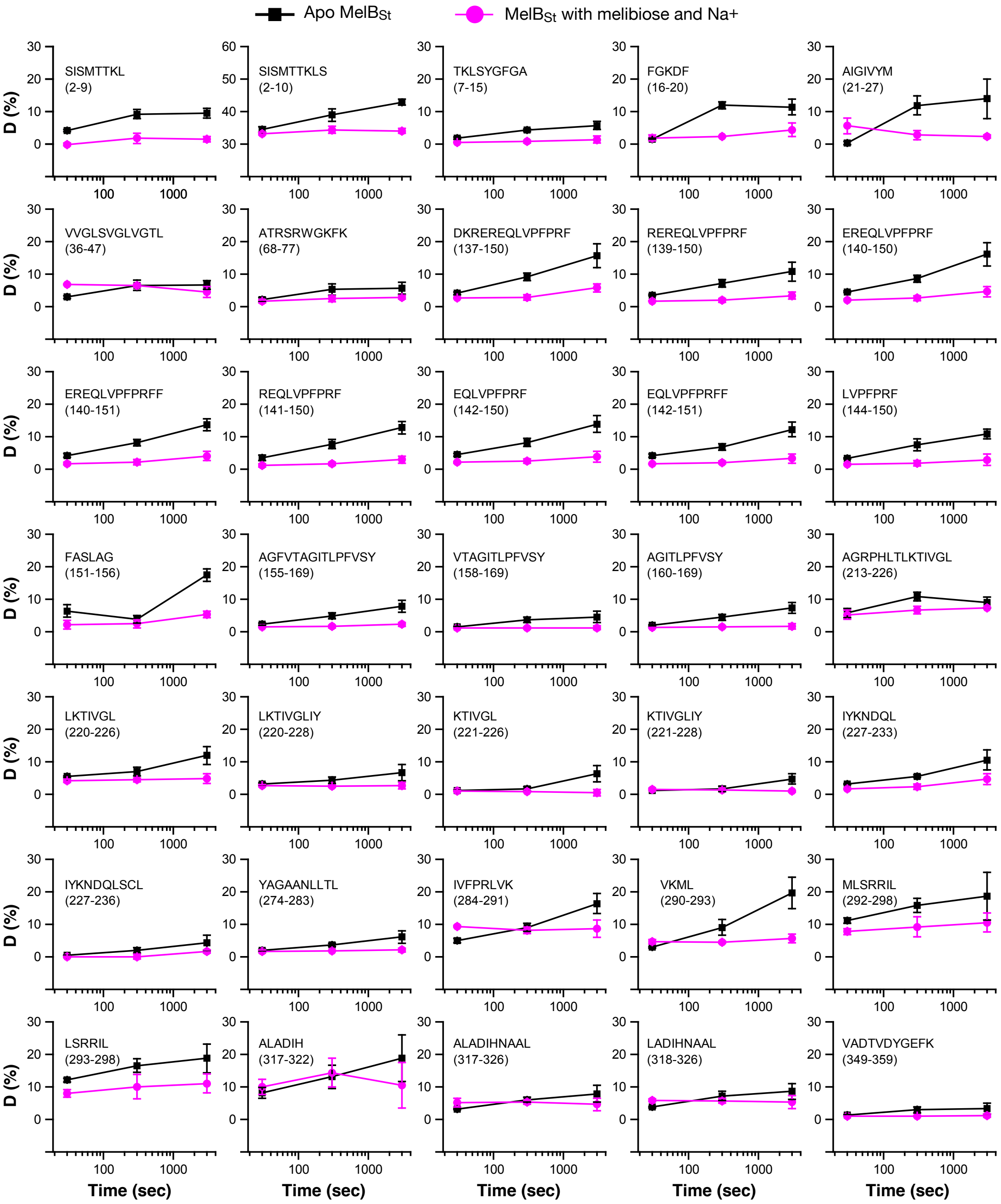

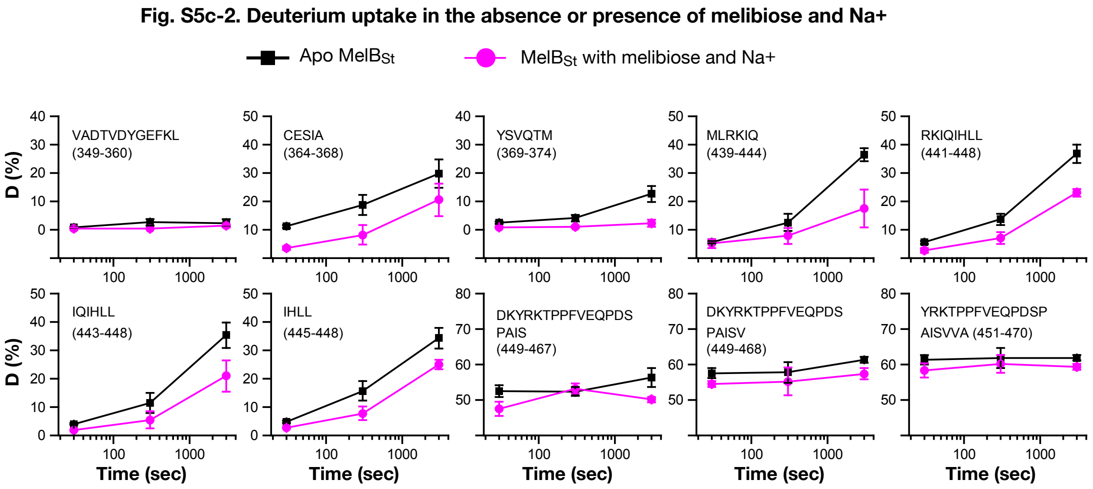


**Fig. S5. Deuterium uptake time course of all peptides ΔD value > threshold.** The percentage of deuterium uptake measured in the absence (filled black square) or presence of melibiose (filled green circle), $Na^+$ (filled red circle), or melibiose and $Na^+$ (filled magenta circle), were plotted against labeling times of 0, 30, 300, and 3000 sec. The peptide sequences and position were shown. **(a)** Melibiose vs. apo. **(b)** $Na^+$ vs. apo. **(c)** Melibiose and $Na^+$ vs. apo. The *y* axe scale was not identical.

Fig. S6

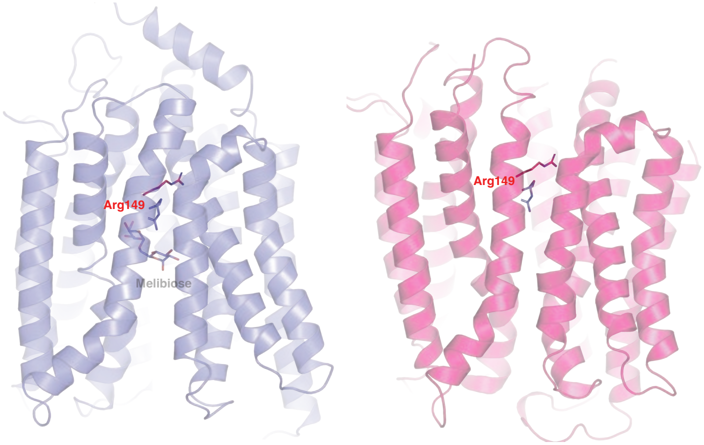


**Outward-facing conformation**

**Inward-facing conformation [PDB 8T60]**

**Fig. S6.** The gating and sugar-binding residue Arg149. Arg149 was shown at the aligned outward- and inward-facing conformations. Arg149 sidechains in red or blue were from the aligned inward-facing [PDB 8T60] and the melibiose-bound outward-facing conformations, respectively.
